# Hepatic ceramide synthesis links systemic inflammation to organelle dysfunction in cancer

**DOI:** 10.1101/2025.10.01.679814

**Authors:** Ying Liu, Ting Miao, Alice Wang, Ezequiel Dantas, Ah-Ram Kim, Zhongjie Zhang, Xiaomei Sun, Richard Binari, John M. Asara, Yanhui Hu, Marcus D. Goncalves, Tobias Janowitz, Norbert Perrimon

**Affiliations:** Department of Genetics, Blavatnik Institute, Harvard Medical School, Boston, MA 02115, USA; Cold Spring Harbor Laboratory, Cold Spring Harbor, New York, NY 11724 USA; Department of Medicine, New York University Grossman School of Medicine, New York, NY 10016, USA; Department of Medicine, Harvard Medical School, Boston, MA, USA; Division of Signal Transduction, Beth Israel Deaconess Medical Center, Boston; Northwell Health Cancer Institute, Northwell Health, New Hyde Park, New York, NY 11042 USA; Howard Hughes Medical Institute, Boston, MA, USA

## Abstract

Paraneoplastic syndromes arise when tumor-derived cytokines reprogram distant organs. Although mediators such as Interleukin-6 have been implicated, how these signals impair host organ function remains incompletely defined. Here, we identify a cytokine-lipid axis that drives hepatic autophagy dysfunction. Specifically, in *Drosophila*, the gut tumor-derived interleukin-like cytokine Upd3 induces the expression of the triglyceride lipase *CG5SCC*, which we named “*cancer-associated lipid mobilizer* (*calm*)”, and the ceramide synthase *schlank* in the fat body. This upregulation rewires fat body lipid metabolism, resulting in an autophagic-flux blockade. Genetic reduction of either *CG5966* or *schlank* restores organelle homeostasis and mitigates paraneoplastic phenotypes. This mechanism is conserved in mammals: in mice, IL-6 upregulates the lipoprotein lipase *Lpl* and ceramide synthases which in turn trigger a hepatic autophagy-flux blockade; in humans, hepatic *LPL* and ceramide synthases expression correlates with poorer survival in hepatocellular carcinoma. Our findings position hepatic lipid metabolism rewiring, especially ceramide synthesis as a critical, conserved node coupling systemic inflammation to organelle dysfunction, and suggest this pathway as a possible therapeutic entry point for cancer-associated liver disorders.

## Introduction

Paraneoplastic syndromes are systemic disorders in cancer patients caused by tumor-derived factors that disrupt distant organs (Pelosof and Gerber, 2010). Among these factors, cytokines, particularly interleukin-6 (IL-6), are prominent paraneoplastic factors that drive diverse clinical manifestations (Ferrer et al., 2023; Flint et al., 2016; Pelosof and Gerber, 2010; Wu et al., 2017). IL-6 engages the JAK/STAT signaling cascade in distant tissues, leading to transcriptional reprogramming that drives systemic inflammation, enhanced thrombopoiesis, and cachectic wasting (Ding et al., 2021; Liu et al., 2025; Stone et al., 2012). One of the primary target tissues of IL-6 in paraneoplastic syndrome is the liver, a central hub of systemic metabolic regulation (Fearon et al., 2012; Liu et al., 2025). In response to chronic IL-6-driven inflammatory cues, the liver enhances gluconeogenesis while promoting adipose lipolysis and skeletal muscle proteolysis, metabolic adaptations that collectively culminate in cancer cachexia. (Ding et al., 2021; Liu et al., 2025). Within hepatocytes, basal autophagy is essential for proteostasis and organelle quality control (Qian et al., 2021). Consistent with this, genetic ablation of core autophagy genes leads to the accumulation of ubiquitinated proteins and p62/SQSTM1-positive inclusions, oxidative stress, hepatomegaly, and hepatic steatosis, underscoring the role of autophagy in maintaining liver function (Komatsu et al., 2007, 2005; Ni et al., 2014). However, whether tumor-derived signals impair hepatic autophagy in a way that contributes to paraneoplastic syndromes remains unknown.

Autophagy is a dynamic membrane-trafficking process that is intrinsically linked to lipid metabolism, making it a highly lipid-demanding pathway. Autophagosome formation begins with the nucleation of an isolation membrane (phagophore), followed by elongation around cytosolic cargo and sealing into a mature vesicle (Ryter et al., 2013; Ueno and Komatsu, 2017). Phospholipids furnish the structural material for phagophore nucleation, membrane elongation, and remodeling during autophagosome biogenesis (Harayama and Riezman, 2018; Rowland et al., 2023). The supply of phospholipids, synthesized from fatty acyl-CoA and glycerol-3-phosphate, sets a key rate constraint; fatty acyl-CoAs are derived from *de novo* lipogenesis or liberated via lipolysis, directly linking autophagic dynamics to cellular lipid metabolism (Harayama and Riezman, 2018; Rowland et al., 2023). Consistently, long-chain acyl-CoA synthetases (ACSLs), which channel fatty acids into *de novo* phospholipid synthesis, relocalize to nascent phagophores to boost local lipid production required for membrane expansion (Dong et al., 2018; Schütter and Graef, 2020). Autophagosome expansion also requires the lipidation and membrane association of ubiquitin-like protein Atg8/LC3: increasing Atg8 enlarges autophagosomes through enhanced elongation and cargo capture, whereas reducing it yields smaller vesicles (Weidberg et al., 2011; Xie et al., 2008). Thus, in metabolically stressed liver, tumor-induced shifts in lipid supply and/or Atg8 abundance are poised to modulate autophagosome number and size, with direct consequences for autophagic capacity (Rowland et al., 2023; Schütter and Graef, 2020; Ueno and Komatsu, 2017).

Dysregulated sphingolipid metabolism exemplifies how lipid imbalances can disrupt autophagy. Ceramides, synthesized *de novo* from serine and palmitoyl-CoA by a family of ceramide synthases (CerS1–6), accumulate in metabolic diseases and some cancers (Hannun and Obeid, 2018; Mullen et al., 2012). While modest increases in ceramide can induce protective autophagy (Taniguchi et al., 2012), chronic or excessive ceramide accumulation impairs autophagic flux, often by compromising lysosomal integrity (Gabandé-Rodríguez et al., 2014; Liu et al., 2016). Mechanistically, sustained ceramide accumulation destabilizes lysosomes, leading to defective hydrolase maturation and leakage of cathepsins into the cytosol, thereby blocking the final stages of autophagic degradation (Gabandé-Rodríguez et al., 2014; Liu et al., 2016; Turk et al., 2001). These findings raise the possibility that tumor-driven perturbation in lipid flux, particularly the buildup of ceramides, could blunt hepatic autophagy and thereby contribute to hepatic impairment and broader systemic dysfunction.

*Drosophila melanogaster* offers a powerful genetic system for dissecting such cross-organ interactions. Multiple fly tumor models exhibit paraneoplastic-like phenotypes, enabling systematic analysis of underlying tumor-host signaling (Bilder et al., 2021; Cheng et al., 2025; Liu et al., 2022). In the Yorkie (Yki) model, gut tumors induced by overexpressing an active form of the oncoprotein Yki, the fly homolog of YAP (*esg»yki^act^*; referred to as Yki flies), elicit systemic effects reminiscent of human paraneoplastic syndromes (Kwon et al., 2015). Yki-driven tumors secrete circulating factors, including PDGF- and VEGF-related factor 1 (Pvf1) and the IL-6-like cytokine Unpaired 3 (Upd3), which act on distant tissues like muscle and fat body to induce wasting (Ding et al., 2021; Liu et al., 2025; Song et al., 2019). The genetic and physiological tractability of the fly fat body, an organ analogous to the vertebrate liver and adipose tissue, makes it an ideal model to investigate how tumor-derived cytokines reshape hepatic lipid metabolism and autophagy to drive paraneoplastic pathology (Liu et al., 2025). Here, guided by a cross-organ gene expression analysis of paraneoplastic progression, we demonstrate that tumor-derived Upd3 activates JAK-STAT signaling to upregulate the lipase *CG5966* and the ceramide synthase *schlank* in the fat body, disrupting lipid homeostasis and triggering severe autophagy dysfunction. Importantly, this pathogenic axis is conserved in mammals, as IL-6 drives expression of lipoprotein lipase (*LPL*) and ceramide synthases in mouse models. Further, elevated *LPL* and *CERS5/6* expression are significantly associated with reduced survival in hepatocellular carcinoma. Together, these findings suggest that hepatic lipid metabolism, namely ceramide synthesis may be a therapeutic target for cancer-associated hepatic dysfunction.

## Results

### Identification of CG5966 as a paraneoplastic factor

In cancer patients, the timing of paraneoplastic syndrome onset frequently diverges from tumor growth (Pelosof and Gerber, 2010). Similarly, in the Yki gut-tumor model, peripheral organ dysfunction (renal failure, muscle degeneration, ovary atrophy) emerges with a delay, robust by day 8 following tumor maturation at day 5 (Kwon et al., 2015; Liu et al., 2025; Song et al., 2019; Xu et al., 2024). To distinguish the effects of tumor burden from tumor-driven systemic signals, we leveraged our full-body single-nucleus transcriptomic data generated previously (Liu et al., 2025), which profiles gene expression dynamics of both gut tumors and host organs at days 5 and 8. We integrated, across cell clusters, both effect size (fold-change) and statistical significance to compute a progression association index for each gene (Fig. 1a). Genes whose differential expression increased from day 5 to day 8 (positive index) were classified as progression-associated and thus prioritized as candidates contributing to paraneoplastic syndrome progression (Fig. 1a). Among cell clusters, enterocytes (ECs) exhibited the largest share of genes associated with progression, consistent with continued differentiation of tumor intestinal stem cells (ISCs) into ECs, followed by follicle cells and indirect flight muscle, reflecting ovarian atrophy and muscle decline, respectively (Fig. S1a) (Liu et al., 2025). Notably, while fat body cell proportions remained relatively stable at both day 5 and day 8 (Liu et al., 2025), it exhibited a higher fraction of progression-associated genes compared with other tissues (Fig. S1a). Top genes identified by the progression association index, including *CG14275, Pepck1, IM2, CG5966,* and *ScsβA*, were predominantly upregulated in the fat body (Fig. 1a; FigS1b-f). Of these, we previously implicated *Pepck1* and *ScsβA* as causal mediators of hepatic gluconeogenic dysregulation and cachectic progression (Liu et al., 2025). Together, these analyses identify the fat body, a fly tissue with many liver-like functions, as a central driver of paraneoplastic progression.

**Figure 1.**
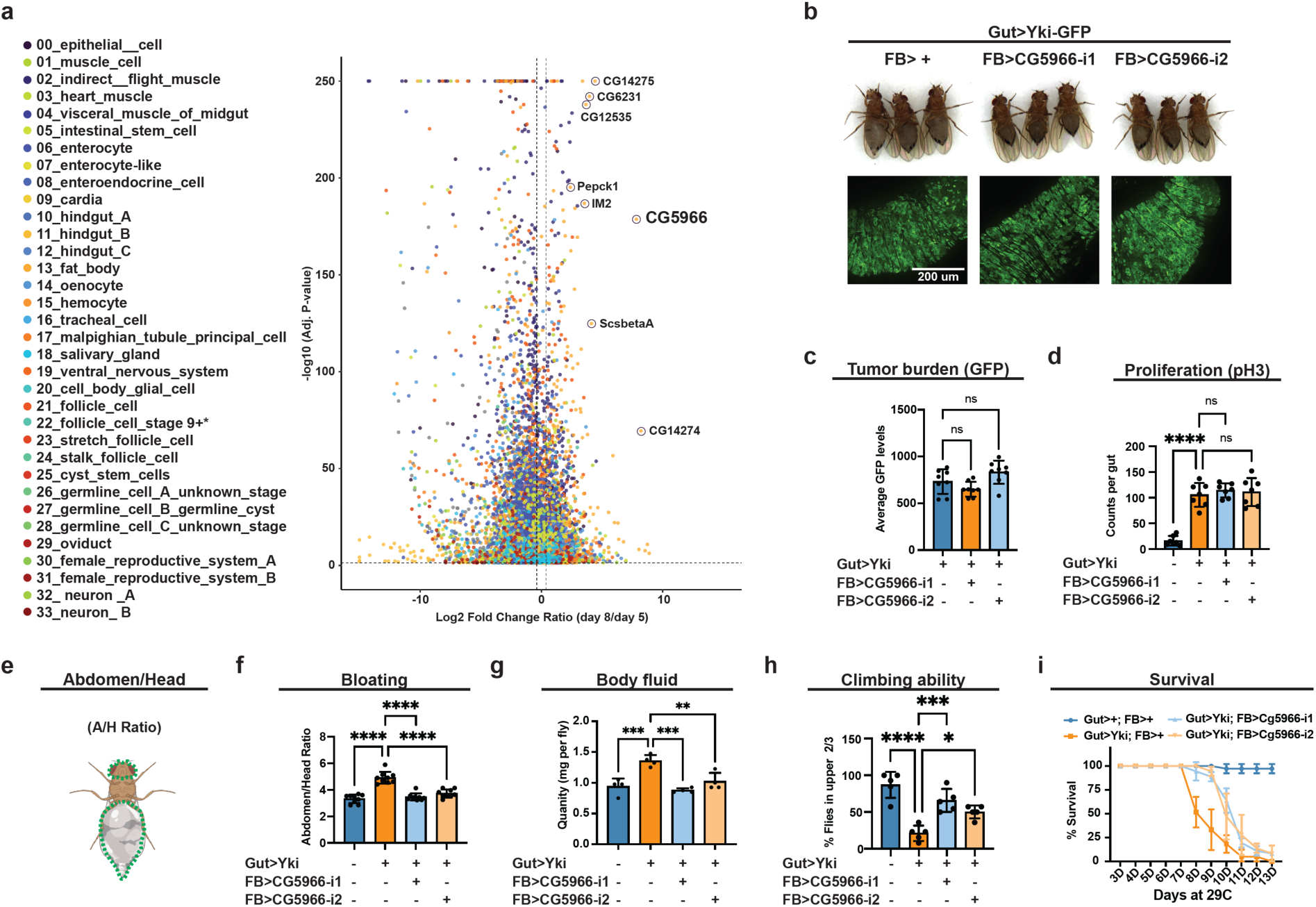
Identification of CG5G66 as a paraneoplastic factor. **a**, Progression association index of differential expressed genes across cell clusters. **b**, Images of gut tumors and phenotypes of Yki flies with or without fat-body *CG5SCC* depletion at day 6. **c**, Quantification of gut tumors (GFP signal, arbitrary units) of Yki flies with or without fat-body *CG5SCC* depletion at day 6 (n = 8). **d**, Total pH3 counts per gut of control flies and Yki flies with or without fat-body *CG5SCC* depletion at day 6 (n = 8). **e**, The bloating quantification by abdomen/head (A/H) size ratio. **f**, Quantification of bloating phenotypes of control flies and Yki flies with or without fat-body *CG5SCC* depletion at day 6 (A/H ratio, n=9). **g**, Body-fluid levels of Yki flies with or without fat-body *CG5SCC* depletion at day 6 (n = 4). **h**,**i**, The climbing ability at day 6 (**h**) (n = 4) and survival curve (**i**) (n = 3) of control flies, Yki flies, and Yki flies with fat-body *CG5SCC* depletion. *p < 0.05, **p < 0.01, ***p < 0.001, ****p < 0.0001, ns indicates not significant. Error bars indicate SDs. n indicates the number of biological replicates in each experiment.

Among the genes with the highest progression association index score, *CG5SCC* is an unannotated gene (Fig. 1a). To test whether its induction contributes to pathogenesis rather than simply reflecting end-stage dysfunction, we combined esg-LexA/LexAop to activate Yki in ISCs (*esg-Le”A, Le”Aop-Yki^act^*) with Lpp-Gal4/UAS to knockdown *CG5SCC* specifically in the fat body (*Lpp-Gal4, UAS-CG5966-RNAi*). Fat-body *CG5966* depletion did not alter GFP-labelled gut tumor size nor phospho-Histone 3 (pH3) signal, a cell mitoses marker, indicating no effect on tumor burden or proliferation (Fig. 1b-d). In contrast, *CG5SCC* knockdown significantly reduced bloating, a hallmark of paraneoplastic phenotype in Yki flies (Kwon et al., 2015; Liu et al., 2025), as evidence by reduced abdomen-to-head (A/H) ratio and total body fluid, suggesting an attenuation of paraneoplastic progression (Fig. 1b, e-g). Consistent with rescue of systemic deterioration, climbing ability improved and lifespan was extended in Yki flies upon fat-body *CG5966* knockdown (Fig. 1h, i). Collectively, these data establish *CG5SCC* as a fat-body-expressed contributor to paraneoplastic pathology in the Yki model.

### CG5G66 targets a subset of triglycerides and reroutes fatty acids into phospholipids

We next investigated the pathogenic mechanism of CG5966. FlyBase annotates *CG5SCC* as a putative triglyceride (TG) lipase. As it associates with cancer and is a lipase, we named *CG5SCC* as *Cancer-Associated Lipid Mobilizer* (*calm*). Sequence alignment with classical human lipases revealed the signature motif G-x-S-x-G in CG5966, as well as the conserved catalytic triad residues (Ser199-Asp227-His312) required for lipid hydrolysis (Fig. S2a). Complementing this, 3D structural modeling using AlphaFold revealed the closest similarity to human pancreatic lipase (PNLIP) and lipoprotein lipase (LPL) (Fig. 2a, b), supporting a close functional relationship between CG5966 and these lipases. Because Yki flies show reduced TG levels (Kwon et al., 2015), we initially hypothesized that upregulated CG5966 simply drives bulk TG hydrolysis to deplete energy stores. However, fat-body-specific *CG5966* knockdown in Yki flies did not restore total TG levels in the abdomen carcass mainly containing fat body tissue (Fig. S2b), suggesting its selective role in TG hydrolysis. To test substrate preference of CG5966 independent of tumors, we profiled lipids in abdomen in wild-type flies with or without fat-body *CG5966* overexpression (Fig. S2c). Consistent with the observations in Yki flies, total TG abundance remained unchanged upon *CG5966* overexpression (Fig. 2c). By contrast, species-resolved lipidomics revealed selective depletion of TGs bearing 16:0, 16:1, 18:0, or 18:1 acyl chain upon *CG5966* overexpression (Fig. 2d-g; S2d), suggesting that CG5966 preferentially acts on specific molecular TGs. To further assess the lipase activity of CG5966, we performed molecular docking using TG (16:0/18:1/18:1), one abundant TG species in both fly and mammals that was significantly depleted upon *CG5966* overexpression. The Boltz-2 docking model positioned the ester bond of the TG substrate in close proximity to the catalytic triad, with a predicted binding free energy of -8.2 kcal/mol, indicating a highly efficient substrate recognition (Fig. 2h) (Mirdita et al., 2022; Passaro et al., 2025). Altogether, our data suggest CG5966 preferentially targets TG containing C16 and C18 acyl chains.

**Figure 2.**
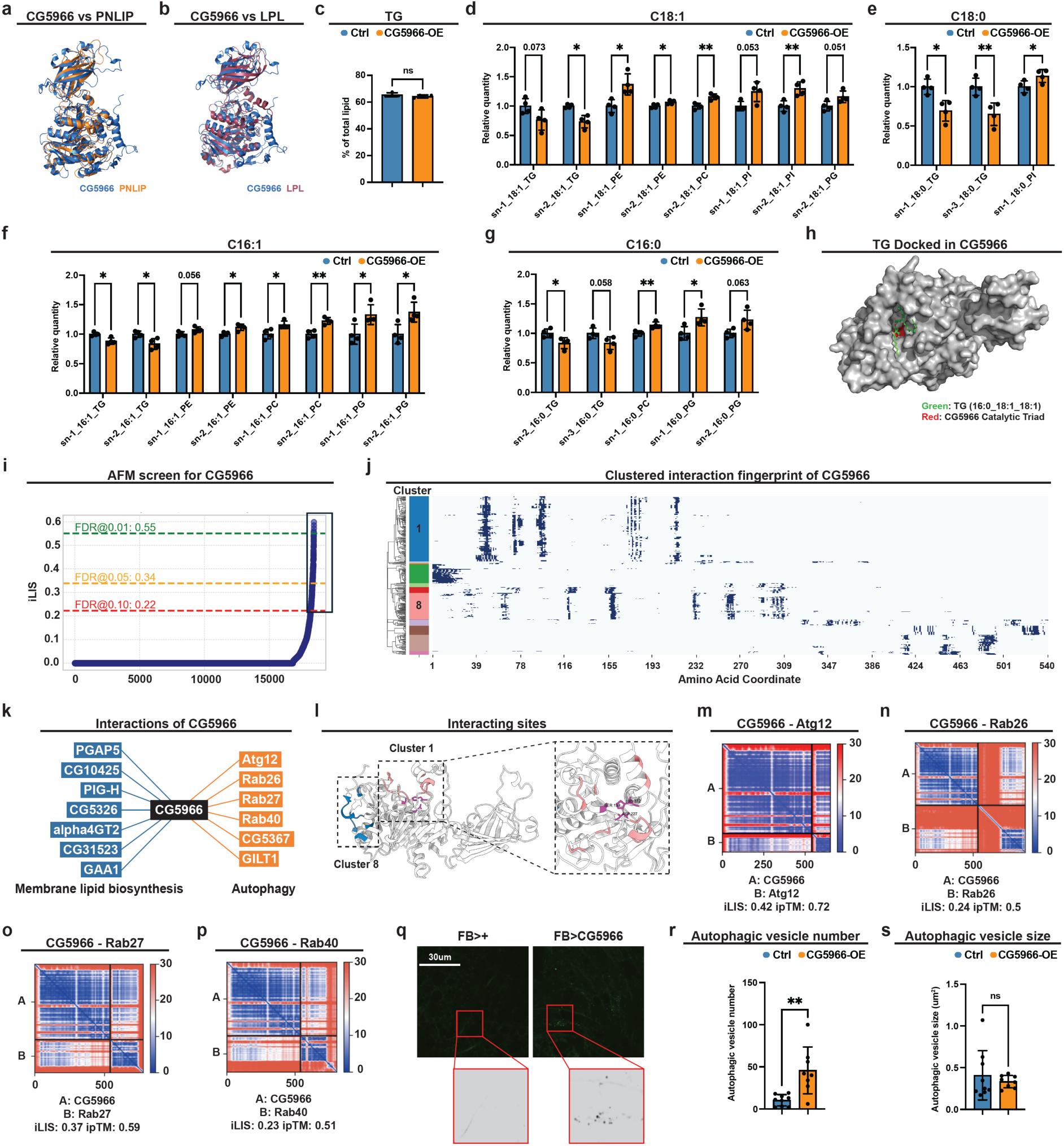
CG5G66 is a triglyceride lipase that promotes autophagosome formation. **a**,**b**, 3D structural modeling using AlphaFold demonstrating similarity of human PNLIP (**a**) and LPL (**b**) to CG5966. **c**, Relative levels of Abdomen TG levels of flies with or without fat-body *CG5SCC* overexpression (n=4), data were retrieved from lipidomics analysis. **d**-**g**, Relative levels of abdomen lipid species with C18:1 (**d**), C18:0 (**e**), C16:1 (**f**), and C16:0 (**g**) acyl chains of flies with or without fat-body *CG5SCC* overexpression (n=4), data were retrieved from lipidomics analysis. **h**, Molecular docking of TG (16:0/18:1/18:1) to CG5966 by the Boltz-2 docking model. **i**, AFM prediction of interacting partners of CG5966. **j**, Cluster of predicted interacting partners of CG5966 based on interacting sites. Y-axis present grouping of interacting patterners and X-axis indicate the interacting sites of CG5966. **k**, Predicted interactions between CG5966 with membrane lipid biosynthesis and autophagy regulators by AFM. **l**, Predicted interacting pocket of CG5966. **m**-**p**, AFM prediction of interaction between CG5966 and Atg12 (**m**), Rab26 (**n**), Rab27 (**o**), Rab40 (**p**). **q**, Live imaging of fat bodies with autophagic vesicle staining dye in flies with or without fat-body *CG5SCC* overexpression at day 7. **r**, **s**, Quantification of autophagic vesicle numbers (**r**) and sizes (**s**) in **q** (n>8). *p < 0.05, **p < 0.01, ***p < 0.001, ****p < 0.0001, ns indicates not significant. Error bars indicate SDs. n indicates the number of biological replicates in each experiment.

As the induction of *CG5966* may not be the leading reason of storage lipid depletion in our tumor model, we investigated the possible physiological consequences of CG5966-mediated C16/18 TG liberation. Lipidomic profiling revealed that phospholipids bearing the same acyl chains were enriched upon *CG5SCC* overexpression: C18:1 increased in Phosphatidylethanolamine (PE), Phosphatidylcholine (PC), Phosphatidylinositol (PI), and Phosphatidylglycerol (PG) (Fig. 2d; Fig. S2e), C18:0 increased in PI (Fig. 2e; Fig. S2f), and C16:1/C16:0 increased in PE/PC/PG (Fig. 2f,g; Fig. S2g,h). As most phospholipids contain C16 and C18 acyl chains (Skotland and Sandvig, 2019), these compositional shifts were companied by increases in total PC, PG, and PI pools (Fig. S2i-k). These data suggest that C16/C18 fatty acyl-CoAs liberated from CG5966-mediated TG depletion are subsequently utilized for phospholipid synthesis. Given that phospholipids are major membrane building blocks (Harayama and Riezman, 2018), we hypothesized that CG5966 play a role in supplying membrane-lipid precursors. If so, it would interact with membrane-lipid biogenesis regulators or enzymes. To test this, we used AlphaFold-Multimer (AFM) to screen putative CG5966 protein-protein interfaces with markers of specific compartments (lysosome, ER, Golgi, plasma membrane) and lipid-metabolic enzymes (Fig. S2l) (Evans et al., 2021). We modeled 3,686 proteins (five AF-M models each; 18,430 total predictions) and, using integrated Local Interaction Score (iLIS) (Kim et al., 2024), retained 248 high-confidence interfaces (1.35%), corresponding to 158 candidate interactors (Fig. 2i; Fig. S2l). Clustering of these candidates by interacting sites yielded 13 groups (Fig. 2j). The most enriched group (Cluster 1) contains membrane-lipid biogenesis regulators, and the second enriched group (cluster 8) includes factors acting at multiple steps of autophagy (Fig. 2j, k). We further checked the interacting interfaces by these top two cluster of proteins, autophagy factors interact CG5966 at out surface, whereas the membrane-lipid biogenesis regulators interact closely to the catalytic triad residues (Ser199-Asp227-His312), suggesting a coordinated procession of lipid (Fig. 2i). Top-scoring partners in the membrane-biogenesis cluster included PGAP5, PIG-H, GPAA1/GAA1 (GPI-anchor synthesis/remodeling) (Kinoshita et al., 2006; Kinoshita, 2020; Tiede et al., 2000), and glycosyltransferases (e.g., α4GT2) implicated in ER/Golgi membrane maturation (Fig. 2k) (Hamel et al., 2010). From the autophagy enriched cluster 8, we recovered autophagy-trafficking factors involved in multiple steps of autophagy process, including Atg12 (ubiquitin-like conjugation system required for autophagosome formation) (Mulakkal et al., 2014), Rab26/27/40 (small GTPases coordinating vesicle dynamics) (Gillingham et al., 2014), CG5367 (Cathepsin L4), and GILT1 (a lysosomal thiol-reductase linked to autolysosome function) (Kongton et al., 2014) (Fig. 2k, m-p). Based on these predictions, we investigated whether CG5966 mediated TG hydrolysis regulates autophagy in the fat body. We assayed autophagy *in vivo* with CYTO-ID® Autophagy Detection Reagent that stains autophagic vesicles (pre-autophagosomes, autophagosomes, and autolysosomes). Indeed, fat-body *CG5SCC* overexpression increased the number of autophagic vesicles, while vesicle sizes were unchanged (Fig. 2q-s). This is consistent with previous report that autophagosome formation depends on the membrane delivery, while size scaling requires other regulators such as ubiquitin-like protein Atg8/LC3 (Weidberg et al., 2011; Xie et al., 2008). Altogether, our results suggest that CG5966 preferentially liberates C16/C18 acyl chains from TG, thereby fueling phospholipid synthesis, expanding membrane lipid supply, and promoting autophagosome formation.

### CG5G66 promotes autophagosome formation in Yki flies

We next asked whether CG5966 modulates autophagy in the Yki model. Fat-body staining in Yki flies revealed a robust increase in number of autophagic vesicles, mirroring the effect of *CG5SCC* overexpression in wild-type animals, with an additional enlargement of vesicle sizes unique to the Yki background (Fig. 3a-c). Both phenotypes (increased vesicle numbers and sizes) were suppressed by fat-body-specific *CG5966* depletion (Fig. 3a-c). These observations suggest a *CG5SCC*-mediated upregulation of autophagosome formation in Yki flies. Elevated autophagosome formation typically reflects enhanced initiation of the autophagy process (Fig. S3a-d). Supporting this, expression of the autophagy-initiating kinase *Atg1* and *Atg17* was elevated in the fat body of Yki flies at day 8 (Fig. S3a-c). To directly test whether autophagy initiation was promoted in Yki flies, we depleted *Atg1* specifically in the fat body of Yki flies. As expected, *Atg1* knockdown blunted autophagic-vesicle staining, restoring vesicle numbers and sizes to normal levels (Fig. 3d-f). This result confirms enhanced autophagy initiation in Yki flies. We then examined whether this activated autophagy initiation contributes to paraneoplastic syndrome. Indeed, fat-body-specific *Atg1* depletion did not alter tumor burden or proliferation (Fig. 3g-i) but significantly reduced bloating and total body fluid (Fig. 3g, j, k) and improved climbing ability (Fig. 3l). Together, these data demonstrate that activation of autophagy initiation contributes to the paraneoplastic syndrome in *Yki* flies, likely mediated by *CG5SCC* through promoting autophagosome formation. Notably, although not observed in *CG5SCC*-overexpressing wild-type flies (Fig. 2q, s), autophagic vesicle enlargement was evident in *Yki* flies and was rescued by either *CG5SCC* or *Atg1* knockdown (Fig. 3a, c, d, f). This discrepancy suggests that in the tumor context, autophagy perturbation extend beyond initiation, implicating additional mediators besides CG5966 in the dysregulation of autophagy in Yki flies.

**Figure 3.**
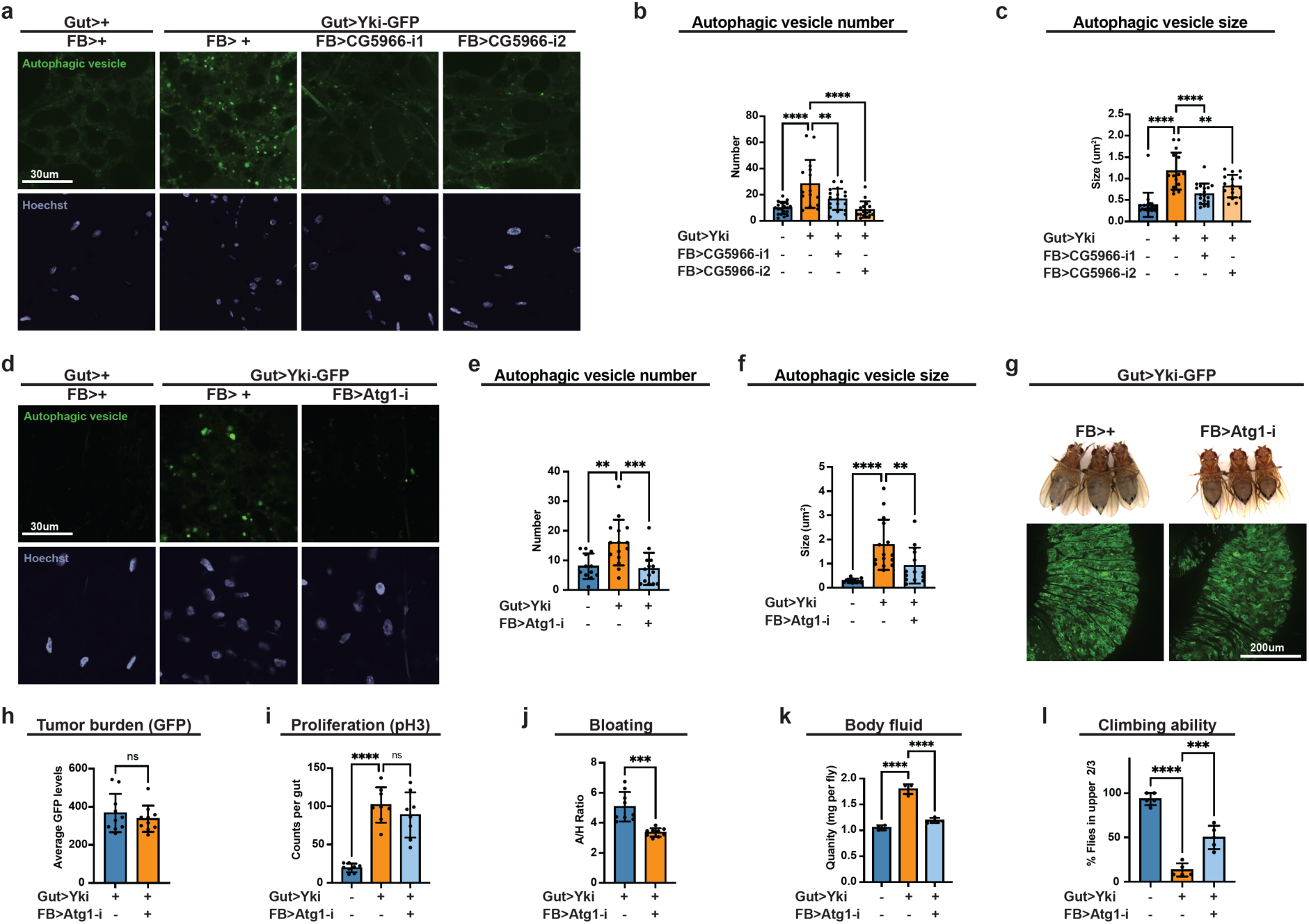
CG5G66 promotes autophagosome formation in the fat body of Yki flies. **a**, Live imaging of fat bodies with autophagic vesicle staining dye in Yki flies with or without fat-body *CG5SCC* depletion at day 7. **b**, **c**, Quantification of autophagic vesicle numbers (**b**) and sizes (**c**) in **a** (n>15). **d**, Live imaging of fat bodies with autophagic vesicle staining dye in Yki flies with or without fat-body *Atg1* depletion at day 7. **e**, **f**, Quantification of autophagic vesicle numbers (**e**) and sizes (**f**) in **d** (n>15). **g**, Images of gut tumors and phenotypes of Yki flies with or without fat-body *Atg1* depletion at day 6. **h**, Quantification of gut tumors (GFP signal, arbitrary units) of Yki flies with or without fat-body *Atg1* depletion at day 6 (n = 8). **i**, Total pH3 counts per gut of control flies and Yki flies with or without fat-body Atg1 depletion at day 6 (n = 8). **j**, Quantification of bloating phenotypes of control flies and Yki flies with or without fat-body Atg1 depletion at day 6 (A/H ratio, n=9). **k**, Body-fluid levels of Yki flies with or without fat-body *Atg1* depletion at day 6 (n = 4). **l**, The climbing ability at day 6 (n = 4) of control flies, Yki flies, and Yki flies with fat-body *Atg1* depletion. *p < 0.05, **p < 0.01, ***p < 0.001, ****p < 0.0001, ns indicates not significant. Error bars indicate SDs. n indicates the number of biological replicates in each experiment.

### Autophagic flux is blocked in Yki flies

We therefore examined classical markers of autophagic flux to clarify the dysregulation in Yki flies. Consistent with the increased number of autophagic vesicles, immunoblotting showed elevated levels of Atg8-II (LC3-II) (Fig. S4a), the lipidated, membrane-bound form of Atg8 associated with autophagosomes (Xie et al., 2008). Because accumulation of Atg8-II can reflect enhanced autophagosome biogenesis and/or flux blockade at autophagosome-lysosome fusion or lysosomal degradation steps, we next assessed autophagic flux and lysosomal function. The anti-p62/Ref(2)P antibody, a negative indicator of autophagic flux, did not yield reliable detection in the adult fat body, so we evaluated cathepsin L (CtsL) maturation as a readout of lysosomal proteolytic activity. CtsL is synthesized as a pro-enzyme that undergoes proteolytic processing into its mature lysosomal form (Turk et al., 2001). Notably, Yki flies displayed elevated levels of pro-CtsL (Fig. 4a), indicating insufficient lysosomal proteolysis. Together with increased autophagosome number and accumulation of Atg8*-*II, these data suggest that autophagosome formation is promoted, whereas lysosomal degradation at the later stage of autophagy is compromised in the fat body of Yki flies. To further assess lysosomal alterations, we examined lysosome morphology and abundance in the fat body of Yki flies using LysoTracker, which labels acidic lysosomes and autolysosomes. Consistent with compromised lysosomal function, Yki fat bodies exhibited abnormally increased size and number of lysosomes, and these alterations were suppressed by fat-body specific *CG5SCC* depletion (Fig. 4b-d). Because a block in lysosomal degradation can arise from defective autophagosome-lysosome fusion or intrinsic lysosomal dysfunction (Fig. S3a), we co-stained autophagic and lysosomal vesicles to examine these possibilities. Co-localization analysis revealed substantial overlap between enlarged autophagic vesicles and lysosomes, indicating that the expanded vesicles in Yki fat bodies are predominantly autolysosomes (autophagosome-lysosome hybrids) (Fig. 4e, f). Notably, the number of autolysosomes were also increased in Yki flies, suggesting no defects in autophagosome-lysosome fusion (Fig. 4g). Consistent with the role of CG5966 in supplying lipids for autophagosome formation, its depletion in Yki flies reduced autolysosome number and was accompanied by smaller autophagic vesicles and lysosomes (Fig. 3c; 4d, g). Together, these results rule out autophagic flux blockade at the autophagosome formation or autophagosome-lysosome fusion steps, instead indicate that the terminal degradation is impaired due to lysosomal dysfunction.

**Figure 4.**
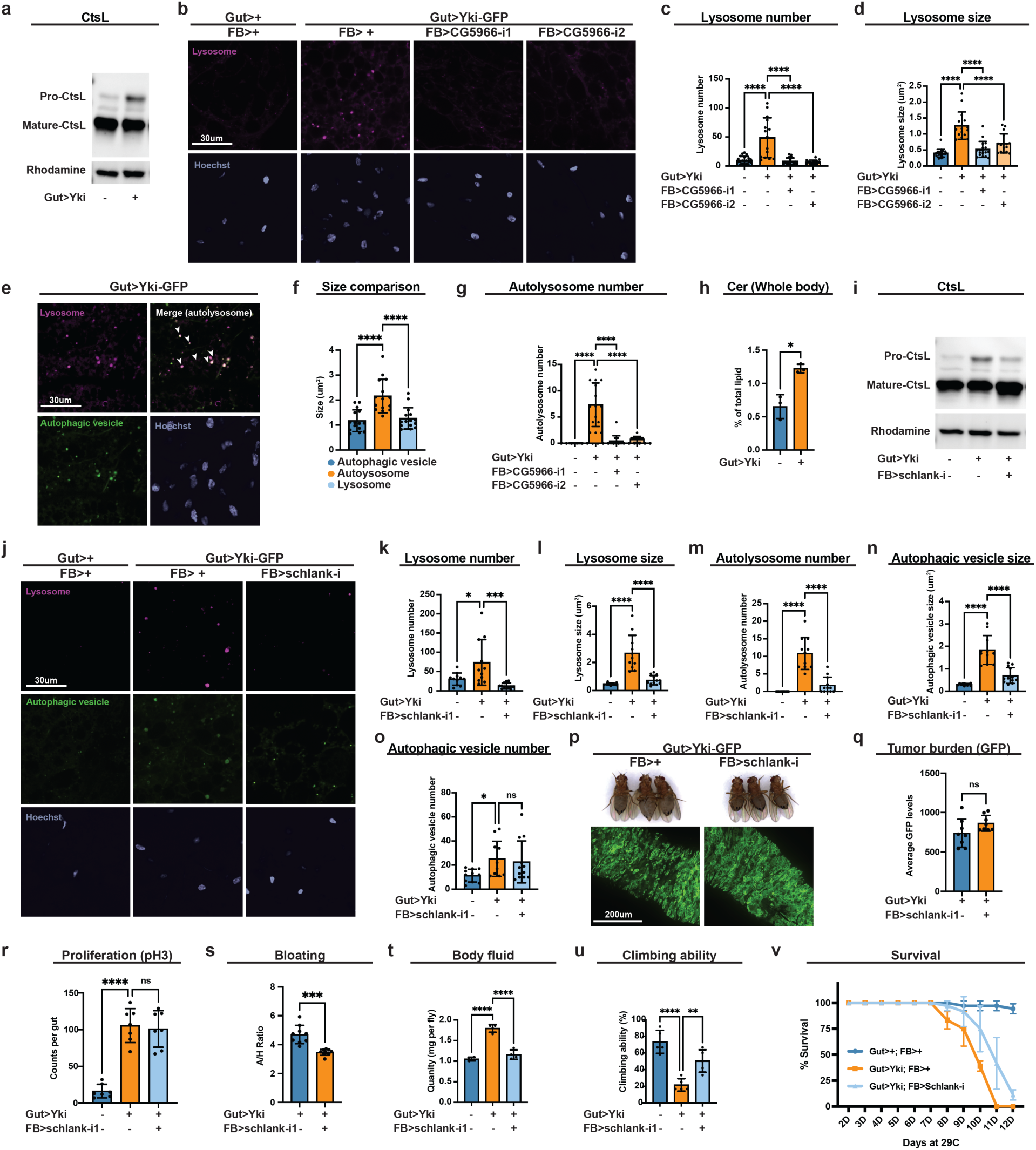
Autophagic flux is blocked in Yki flies. **a**, Western blot showing pro and matured forms of CtsL in Yki and control flies. Rhodamine was used as loading control. **b**, Live imaging of lysosomes in Yki flies with or without fat-body *CG5SCC* depletion at day 7. **c**, **d**, Quantification of lysosome numbers (**c**) and sizes (**d**) in **b** (n>15). **e**, Live imaging of lysosome and autophagic vesicles in Yki flies at day 7. **f**, Quantification of sizes of autophagic vesicles, autolysosome and lysosome in Yki flies (n>15). **g**, Quantification of autolysosome numbers in Yki flies with or without fat-body *CG5SCC* depletion at day 7 (n>15). **h**, Relative levels of whole-body ceramide levels of Yki and control flies (n=3), data were retrieved from lipidomics analysis. **i**, Western blot showing pro and matured forms of CtsL in control flies, Yki flies with or without fat-body *schlank* depletion at day 7. Rhodamine was used as loading control. **j**, Live imaging of lysosomes and autophagic vesicles in Yki flies with or without fat-body *schlank* depletion at day 7. **k**-**n**, Quantification of lysosome numbers (**k**) and sizes (**l**), autophagic vesicle numbers (**m**) and sizes (**n**), autolysosome numbers (**o**) in **j** (n>15). **p**, Images of gut tumors and phenotypes of Yki flies with or without fat-body *schlank* depletion at day 6. **q**, Quantification of gut tumors (GFP signal, arbitrary units) of Yki flies with or without fat-body *schlank* depletion at day 6 (n = 8). **r**, Total pH3 counts per gut of control flies and Yki flies with or without fat-body *schlank* depletion at day 6 (n = 8). **s**, Quantification of bloating phenotypes of control flies and Yki flies with or without fat-body *schlank* depletion at day 6 (A/H ratio, n=9). **t**, Body-fluid levels of Yki flies with or without fat-body *schlank* depletion at day 6 (n = 4). **u**, **v**, The climbing ability at day 6 (**u**) (n = 4) and survival curve (**v**) (n = 3) of control flies, Yki flies, and Yki flies with fat-body *schlank* depletion. *p < 0.05, **p < 0.01, ***p < 0.001, ****p < 0.0001, ns indicates not significant. Error bars indicate SDs. n indicates the number of biological replicates in each experiment.

## Ceramide accumulation underlies lysosomal dysfunction in Yki flies

We next investigated why terminal lysosomal degradation is impaired in the fat body of Yki flies. Intrinsic lysosomal dysfunction can result from several causes, including defective lysosomal acidification and compromised hydrolase processing (Colacurcio and Nixon, 2016; Saftig and Klumperman, 2009). Surveying our Yki transcriptome and lipidomics datasets, we found no evidence of altered expression in genes governing lysosomal acidification (e.g., *Vha100-1*, *Vha16-1*, *VhaAC39-1*) or hydrolase processing (e.g., *Lerp*, *Rab5*, *Rab6*, *Rab7*, *Sy”17*); with one exception: *CtsL1* was upregulated (Fig. S4b). Together with the accumulation of pro-CtsL, this pattern is consistent with a possible compensatory response to impaired CtsL maturation. In contract, we noticed a significant increase in sphingolipids among the major lipid classes (Fig S4c-g), with a double amount of whole-body ceramide levels in Yki flies compared with controls (Fig. 4h). Excessive ceramides are known to disrupt lysosomal function and blocking cargo degradation (Gabandé-Rodríguez et al., 2014; Liu et al., 2016). These findings suggest a ceramide-driven mechanism for the lysosomal defect in Yki flies. To test this hypothesis, we depleted *schlank*, the sole ceramide synthase characterized in *Drosophila*, in the fat body and assessed lysosomal function by CtsL maturation. As predicted, *schlank* knockdown reduced the accumulation of pro-CtsL in Yki flies, indicating improved lysosomal proteolysis (Fig. 4i). Concordantly, *schlank* depletion restored lysosome number and size to near-normal levels (Fig. 4j-l). Importantly, co-staining of autophagic and lysosomal vesicles showed reduced autolysosome number (Fig. 4m), and the enlarged autolysosomes were normalized close to typical autophagic vesicle dimensions (Fig. 4n). Notably, *schlank* knockdown did not change autophagic vesicle numbers (Fig. 4o), consistent with our model that autophagosome biogenesis is mediated by CG5966. Together, these findings indicate that *Schlank* depletion enhances lysosomal clearance and alleviates autophagic blockade in *Yki* flies. We next accessed whether relieving the lysosomal blockade improves host physiology. Without altering tumor burden or proliferation (Fig. 4p-r), *schlank* depletion reduced bloating and total body fluid (Fig. 4p, s, t), restored climbing ability, and extended survival in Yki flies (Fig. 4u, v). Together, these findings support a model in which ceramide accumulation-induced lysosomal dysfunction blocks autophagic clearance; the consequent accumulation of enlarged, dysfunctional autolysosomes contribute causally to paraneoplastic syndrome pathology in the Yki model.

### The JAK–STAT pathway drives fat body autophagy dysregulation

We next asked what upstream signal induces fat-body autophagy dysregulation. Many TG lipases (e.g., *brummer/bmm*, fly homolog of *ATGL*) are upregulated when insulin signaling is low to mobilize stored energy (Wang et al., 2011). Although Yki flies exhibit reduced systemic insulin activity due to elevated circulating ImpL2 (an insulin-like peptide antagonist) (Kwon et al., 2015), *CG5966* expression was not suppressed by restoring fat body insulin signaling via overexpression of a constitutively active from of the insulin receptor (*InRca*) (Fig. S5a). Beyond insulin, prior work showed increased Upd3/JAK-STAT and Pvf1/Pvr-RTK signaling in Yki flies (Ding et al., 2021; Liu et al., 2022, 2025; Song et al., 2019); we therefore tested whether these pathways regulate *CG5SCC*. Among these candidates, *Upd3* overexpression in intestinal stem cells (ISCs) (*esg»Upd3*) led to an upregulation of *CG5SCC* in the fat body (Fig. 5a). Consistently, fat body expression of constitutively active from of Stat92E (*Lpp»StatS2E^act^-HA*) increased *CG5SCC* expression (Fig. 5b). Strikingly, the ceramide synthase *schlank* showed the same regulatory pattern (Fig. 5c, d). These results suggest that the Upd3/JAK-STAT pathway induces expression of both *CG5SCC* and *schlank* in wildtype flies.

**Figure 5.**
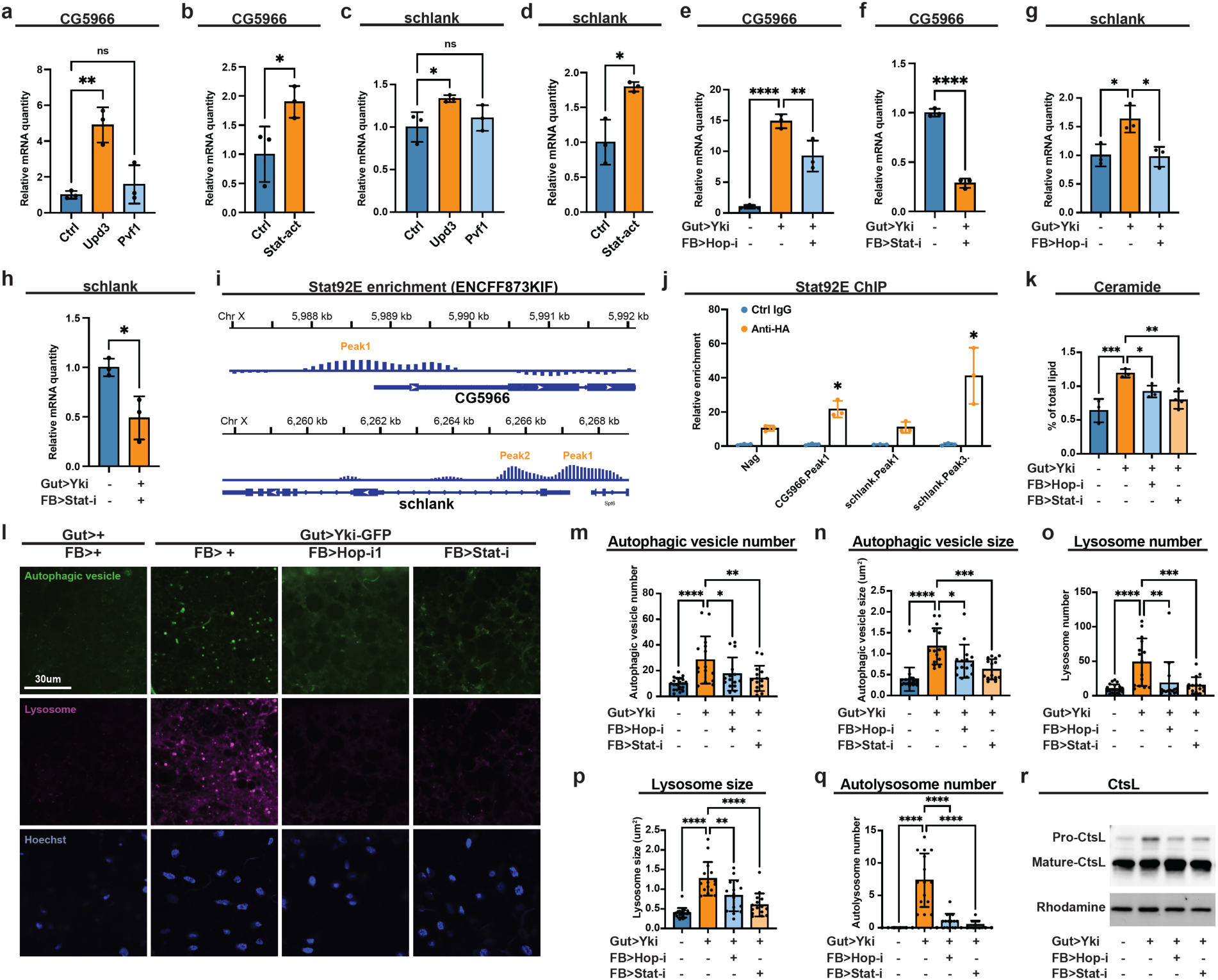
The JAK–STAT pathway regulates *CG53CC* and *schlank* expression in the fat body. **a**, **c**, qRT–PCR of *CG5SCC* (**a**) and *schlank* (**c**) mRNA in the fat body of flies with or without ISC *Upd3* or *Pvf1* expression (*esg*>*Upd3* or *Pvf1*) at day 8 (n = 3). **b**, **d**, *CG5SCC* (**b**) and *schlank* (**d**) mRNA in the fat body of flies with or without fat-body Stat92E-act (constitutive-activated form of Stat92e) expression at day 8 (n = 3). **e**, **g**, *CG5SCC* (**e**) and *schlank* (**g**) mRNA in the fat body of control, Yki flies, and Yki flies with fat-body *hop* depletion at day 6 (n = 3). **f**, **h**, *CG5SCC* (**f**) and *schlank* (**h**) mRNA in the fat body of Yki flies with or without fat-body *StatS2e* depletion at day 6 (n = 3). **i**, Data from the ChIP-seq database indicating enrichment of Stat92e binding at the *CG5SCC* and *schlank* promoter and gene regions. **j**, ChIP revealed the enrichment of HA-tagged Stat92E-act binding at the *CG5SCC* and *schlank* promoter and gene regions, shown by fold changes relative to control IgG at day 8 (n = 3). Neg, negative control. **k**, Relative levels of whole-body ceramide levels of control, Yki flies, and Yki flies with fat-body *hop or StatS2E* depletion at day 6 (n > 3), data were retrieved from lipidomics analysis. **l**, Live imaging of lysosome and autophagic vesicles in Yki flies with or without fat-body *hop* or *StatS2e* depletion at day 7. **m**-**q**, Quantification of autophagic vesicle numbers (**m**) and sizes (**n**), lysosome numbers (**o**) and sizes (**p**), and autolysosome numbers (**q**) in **l** (n > 15). **r**, Western blot showing pro and matured forms of CtsL in control flies, Yki flies with or without fat-body *hop* or *StatS2e* depletion at day 7. Rhodamine was used as loading control. The same membrane was probed sequentially with Atg8 antibody in S4a, loading control is shared. *p < 0.05, **p < 0.01, ***p < 0.001, ****p < 0.0001, ns indicates not significant. Error bars indicate SDs. n indicates the number of biological replicates in each experiment.

To validate this regulatory axis in the tumor context, we used the dual binary systems: LexA-LexAop to express Yki in ISCs and GAL4-UAS to deplete the JAK-STAT pathway components *hop* (JAK) and *Stat92e* (STAT) in the fat body. As expected, inhibition of JAK-STAT pathway reduced *CG5SCC* and *schlank* expression (Fig. 5e-h). Published Stat92e ChIP-seq dataset (ENCODE, ENCSR290OJD/ENCFF873KIF) revealed putative Stat92E occupancy within promoter and intronic regions of *CG5966* and *schlank* (Fig. 5i). In agreement, fat body ChIP confirmed Stat92E association at these loci (Fig. 5j). Consistent with the gene expression regulation, inhibiting JAK-STAT signaling in the fat body lowered whole-body ceramide (Fig. 5k), reduced the size and number of autophagic vesicles and lysosomes, decreased autolysosome number (Fig. 5i-q), and cleared the accumulation of pro-CtsL (Fig. 5r), indicating restored lysosomal maturation. Together, these data indicate that Upd3/JAK-STAT signaling upregulates two effectors of autophagy in the fat body: *CG5966*, which boosts autophagosome biogenesis; and *schlank*, which elevates ceramide and impairs lysosomal function, together driving autophagy dysregulation in Yki flies.

### Conserved pathogenic mechanism in mouse models and humans

Public STAT3 ChIP-seq dataset (ENCODE, ENCSR000DEC) in human Hela cells indicates putative STAT3 occupancy within the promoter regions of *LPL* and *CERS5* (Fig. S5b, c), suggesting a conserved regulation in mammals. To test whether this JAK-STAT-mediated lipid-autophagy pathogenic axis is conserved, we employed a mouse IL-6 tumor model: C57BL/6 mice bearing Lewis lung carcinoma (LLC) tumors engineered either to secrete IL-6 (LLC+IL-6) or not (LLC). LLC+IL-6 mice exhibit rapid weight loss, indicating progress of paraneoplastic syndrome (Liu et al., 2025). To determine whether mammalian lipases functionally homologous to CG5966 act downstream of IL-6 in tumor-bearing mice, we profiled the expression of its putative orthologs (*Pnlip* and *Lpl*) in the liver as well as adipose tissue, the major TG storage site in mammals. *Pnlip*, predominantly expressed in the pancreas, was undetectable in adipose tissue or liver, whereas *Lpl* was induced in both tissues of LLC+IL-6 mice (Fig. 6a, b). These data indicate that IL-6 promotes lipolysis in both adipose tissue and liver, aligning with our observation of elevated lipolysis in the fly fat body, which is functionally analogous to these mammalian tissues. To evaluate functional conservation, we performed lipidomic profiling of adipose tissue and liver in LLC and LLC+IL-6 mice. In adipose tissue, TG species containing C16/C18 acyl chains were significantly reduced in LLC+IL-6 mice (Fig. 6c; Fig. S6a), suggesting preferential liberation of C16/18 fatty acids through activated lipolysis. A similar selective TG depletion occurred in liver (Fig. 6d; Fig. S6b), accompanied by significant enrichment of phospholipids (PC, PI, PE, PG) containing C16/C18 chains (Fig. 6e-h; Fig. S6c-e), mirroring our observation in flies. Together, these findings support a conserved IL-6-driven program that mobilizes C16/18 fatty acids from TG to supply specific phospholipid synthesis. Given that lipolysis substrates reside primarily in adipocytes in mammals, these results point to an adipose-liver metabolic circuit in which adipocyte TG hydrolysis provides fatty acids that fuel hepatic phospholipid biogenesis.

**Figure 6.**
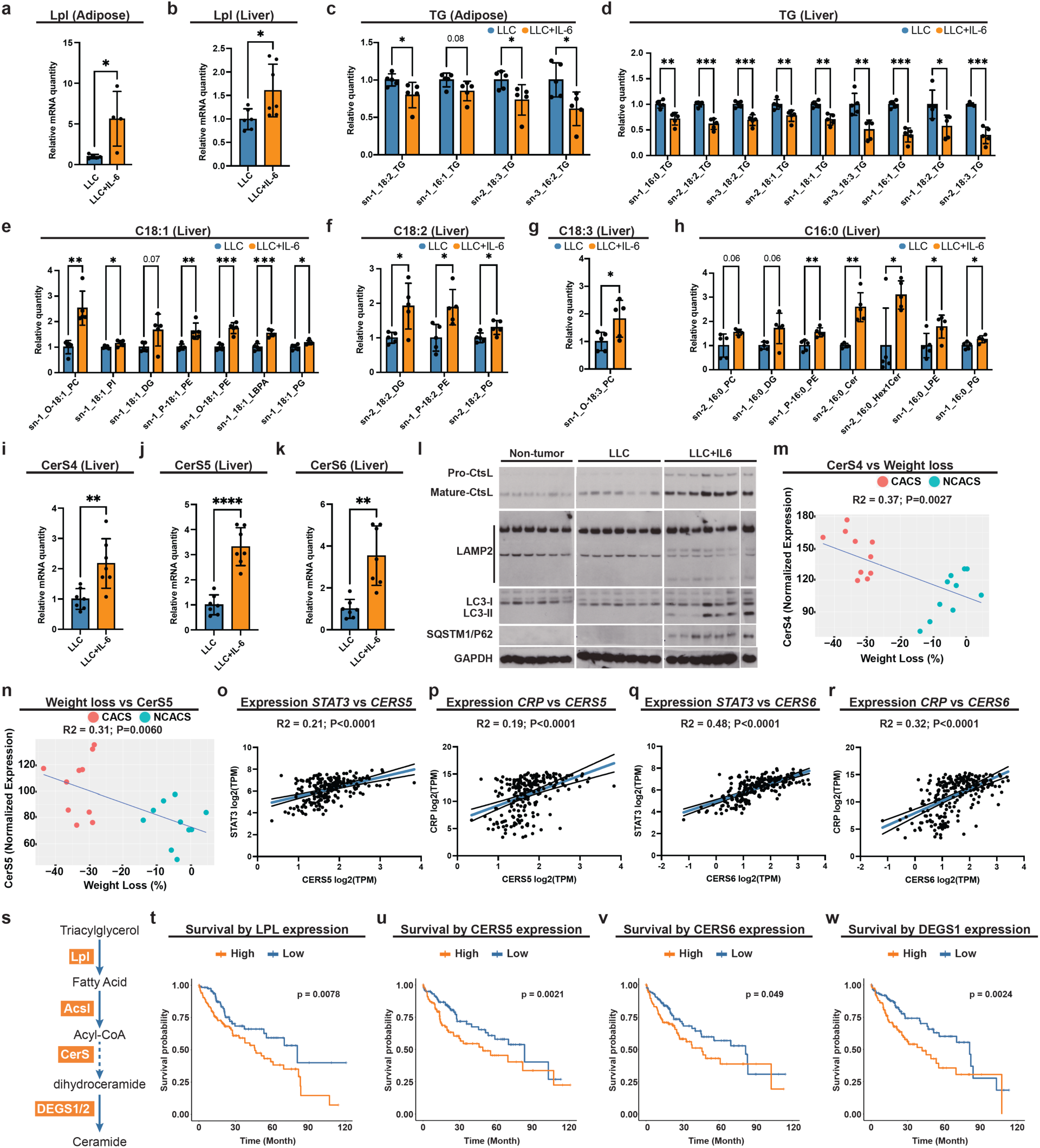
Conserved pathogenic mechanism in mouse models and humans. **a**, **b**, qRT–PCR of *Lpl* mRNA in adipose tissue (**a**) and liver (**b**) B6 mice injected with LLC cells with (n = 4) or without (n = 5) and IL-6 expression. **c**,**d**, Relative levels of adipose (**c**) and liver (**d**) TG species of B6 mice injected with LLC cells with or without IL-6 expression (n=5). **e**-**h**, Relative levels of liver lipid species with C18:1 (**e**), C18:2 (**f**), C18:3 (**g**), and C16:0 (**h**) acyl chains of B6 mice injected with LLC cells with or without IL-6 expression (n=5). **i**-**k**, qRT–PCR of *CerS4* (**i**), *CerS5* (**j**), *CerSC* (**k**) mRNA in liver of B6 mice injected with LLC cells with or without IL-6 expression (n = 7). **i**, Western blot showing forms of CtsL, LAMP2, LC3, and P62 in liver of B6 mice injected with LLC cells with or without IL-6 expression, with the loading control GAPDH. **m**,**n**, Correlation plots showing the positive relationship between liver *CerS4* (**m**) and *CerS5* (**n**) expression (n = 10) versus weight loss of KL mice. **o**-**r**, Correlation plots showing the positive relationship between expression of *STAT3* and *CerS5* (**o**), *CPR* and *CerS5* (**p**), *STAT3* and *CerSC* (**q**), *CPR* and *CerSC* (**r**) in non-diseased liver samples of 226 participants in the GTEx Project. TPM, transcript per million. **s**, Pathway map showing *de novo* ceramide synthesis pathway in human. **t**-**w**, Kaplan– Meier survival curves displaying the estimated survival probabilities of people with hepatocellular carcinoma with low (bottom third) and high (top third) hepatic expression of *LPL* (**t**), *CERS5* (**u**), *CERSC* (**v**), and *DEGS1* (**w**). *p < 0.05, **p < 0.01, ***p < 0.001, ****p < 0.0001, ns indicates not significant. Error bars indicate SDs. n indicates the number of biological replicates in each experiment.

We next examined whether JAK-STAT dependent upregulation of hepatic ceramide synthesis is also conserved in mice. Among the six *CerS* genes (*CerS1-6*), *CerS4*, *CerS5*, and *CerS6* were upregulated in liver following IL-6 induction (Fig. 6i-k; Fig. S6f). Unlike the fly enzyme schlank (with limited substrate bias), mammalian CerS isoforms exhibit distinct acyl-chain specificities (Mizutani et al., 2005; Pewzner-Jung et al., 2010; Riebeling et al., 2003). Concordant with the known C16 preference of CerS5/6 (Mizutani et al., 2005; Pewzner-Jung et al., 2010; Riebeling et al., 2003), we observed a selective increase in C16:0 ceramide in LLC+IL-6 mice (Fig. 6h), supporting targeted ceramide production. Given previous report that hepatic ceramides are synthesized from acyl-CoAs (Pewzner-Jung et al., 2010), together with our observation that hepatic TG species containing C16:0 acyl chains were significantly reduced (Fig. 6d), it is likely that elevated C16 ceramide synthesis in LLC*+*IL*-6* mice is fueled by palmitoyl-CoA derived from enhanced lipolysis.

Next, we asked whether autophagic flux is blocked in mouse tumor model similarly as observed in our fly model. Immunoblot markers of autophagic flux indicated a late-stage block at the autolysosome in livers of LLC+IL-6 mice (Fig. 6l), evidenced by: accumulation of pro-CtsL, indicating impaired lysosomal maturation; elevated SQSTM1/p62, indicating defective cargo degradation; incompletely glycosylated LAMP2, consistent with lysosomal biogenesis/maturation defects; and increased LC3-II, which, in the context of rising p62, supports impaired flux downstream of autophagosome formation (Klionsky et al., 2021). Together, these findings indicate blocked autophagic clearance in the IL6 tumor-bearing mice. To access the paraneoplastic consequences of these autophagy dysregulations, we turned to an inducible lung-cancer model (*Kras^LSL-G12D^/+; Lkb^^ffox/ffox^*; “KL” mice), in which tumors were initiated by intranasal Ad-Cre. In this model, ∼70% of mice display cachexia (>15% body-weight loss) 5-6 weeks following the tumor induction (Goncalves et al., 2018). We assessed the expression correlation of *CerS* and their weight loss in cachectic and non-cachectic groups, and found that expression of *CerS4* and *CerS5*, which preferentially synthesize C18-20 and C16 ceramides, respectively, were positively correlated with weight loss (Fig. 6m, n), reinforcing their association with paraneoplastic syndrome progression.

Extending to humans, we analyzed non-diseased liver from GTEx project (226 participants). Because IL-6 activates STAT3, we asked whether hepatic *CERS4/5/C* expression corresponds with STAT3 activity. Expression of *CERS5* and *CERS6* positively correlated with canonical IL-6/STAT3 targets (*CRP*, *HIF1A*, *TIMP1*) and with *STAT3* itself (Fig. 6o-r; Fig. S6g-j) (Castell et al., 1990; Dien et al., 2006; Johnson et al., 2018; Yu et al., 2007), suggesting IL-6 driven regulation of *CERS5* and *CERS6* in the human liver. To explore the clinical relevance of hepatic lipolysis and ceramide synthesis, we analyzed transcriptomic data from the TCGA Pan-Cancer Hepatocellular Carcinoma cohort (372 participants) (Hoadley et al., 2018; Liu et al., 2018). In alignment with our fly and mouse model findings, higher hepatic *LPL* expression associated with worse survival (Fig. 6t). We then examined genes in human *de novo* ceramide synthesis pathway (Fig. 6s): *CERS5* and *CERS6* were significantly associated with worse survival (Fig. 6u, v; Fig. S6k-n), and higher expression of *DEGS1*, which catalyzes the desaturation of dihydroceramide to ceramide (Fig. 6s), aligned with adverse outcomes (Fig. 6w). By contrast, *DEGS2*, a bifunctional C4-hydroxylase/desaturase with tissue-restricted roles (Ota et al., 2023; UniPort Q6QHC5), showed no such association, consistent with its lower hepatic relevance (Fig. S6o). Collectively, these cross-species data support a conserved mechanism: Upd3/IL-6/JAK-STAT signaling induces *LPL* and *CerS* expression (notably CerS5/6), driving a late-stage autophagy block that contributes to paraneoplastic syndrome.

## Discussion

IL-6 is a central cytokine in the metabolic crosstalk between tumors and the host, coordinating systemic inflammation and altered fuel handling that drive paraneoplastic syndromes (Barton, 2001; Johnson et al., 2018). Our work builds upon previous studies by identifying a novel, cytokine-driven lipid-autophagy axis that underlies paraneoplastic hepatic dysfunction, specifically: (1) by charting paraneoplastic-syndrome associated genes organism-wide, we identify two key drivers: the triglyceride lipase *calm*/*CG5SCC* (putative human *LPL* ortholog) and the ceramide synthase *schlank* (human *CERS* ortholog); (2) we reveal a previously unrecognized Upd3/JAK-STAT control of hepatic lipid metabolism through direct transcriptional upregulation of both genes; (3) we show that this lipid reprogramming initiates autophagosome formation yet blocks autophagic completion, leading to a pathologic buildup of enlarged, dysfunctional autolysosomes and consequently hepatic injury; (4) we validate conservation in preclinical mouse models, where IL-6 elevates hepatic *Lpl* and *CerS* alongside a late-stage autophagy block; (5) we extend these findings to humans, showing that higher hepatic *LPL* and *CERS* expression correlates with poorer survival in hepatocellular carcinoma, thereby nominating the IL-6/LPL-CERS axis as a tractable therapeutic target.

JAK-STAT signaling orchestrates systemic energy mobilization through coordinated actions across tissues. Through STAT3 activation, Leukemia inhibitory factor (LIF) suppresses hepatic lipogenic gene expression and increases adipose triglyceride lipase (ATGL) activity in adipose tissue, thereby promoting FAs release from TG (Arora et al., 2018; Yang et al., 2024). Because IL-6 also activates STAT3, similar repression of lipogenesis and activation of lipolysis via ATGL and/or other lipases are expected with enhanced IL-6 signaling. Indeed, our data show that IL-6 upregulates *LPL* in adipose tissue and liver, which may act with ATGL to promote free FAs release from TG. Under physiological conditions, excess FAs are re-esterified into TAG and phospholipids or oxidized in mitochondria via β-oxidation to feed the TCA cycle (Houten and Wanders, 2010; Lee and Ridgway, 2020). We previously showed that hepatic IL-6/JAK-STAT3 signaling induces gluconeogenic programs and elevates *PDK3*, which inhibits pyruvate dehydrogenase and restricts glycolytic acetyl-CoA entry into the TCA cycle (Liu et al., 2025). Thus, upon activation of JAK-STAT pathway, hepatocytes may shift substrate of TCA cycle from glucose (glycolysis-derived) to fatty acids (β-oxidation-derived). While this may be adaptive during acute stress (Qing et al., 2020), chronic IL-6 signaling imposes sustained FA influx that can exceed hepatic β-oxidation capacity (Kerner et al., 2008), prompting diversion of surplus FAs into alternative lipid pathways, including phospholipid and ceramide synthesis. We therefore propose that IL-6’s normally adaptive substrate-mobilizing role becomes maladaptive in cancer: excess FAs that cannot be efficiently oxidized are progressively redirected into membrane lipids and bioactive sphingolipids, predisposing to lipid-driven organelle dysfunction.

Hepatic lipid metabolic reprogramming is an active driver of paraneoplastic pathology. Although depletion of storage lipid is frequently observed in cancer and often interpreted solely as a signature wasting phenotype accompanied by disease progression, our findings reveal that TG depletion and reprogramming of phospholipid and ceramide metabolism contribute causally to organelle dysfunction, uncovering a novel mechanism contributing to hepatic dysfunction and systemic paraneoplastic disorders. In Yki flies, fatty-acid overload augments the membrane lipids supply required for autophagosome formation. In line with the lipid composition of autophagic structures in flies, where PE species with total carbon 32 and 34 predominate (C16/C16, C16/C18, C14/C18) (Laczkó-Dobos et al., 2021), we show that CG5966 preferentially mobilizes C16 and C18 fatty acids from TG, enriching C16-and C18-containing PE to supply membranes for autophagosome formation. Notably, this heightened initiation is tolerated, as *CG5SCC* overexpression alone in flies does not trigger autophagy dysfunction or induces paraneoplastic phenotypes. By contrast, in the context of Yki, CG5966-mediated membrane lipid supply and concomitant ceramide synthesis is deleterious. Sustained ceramide accumulation compromises lysosomal function, inducing membrane permeabilization, disrupting cathepsin processing, and blocking late-stage autophagic clearance (Gabandé-Rodríguez et al., 2014; Liu et al., 2016). Accordingly, we observe defective lysosomal maturation in Yki flies and in tumor-bearing mice, indicating that cancer-induced lipid reprogramming actively drives autophagy failure and hepatic dysfunction rather than representing a mere by-product of systemic disease.

IL-6/ceramide-induced lysosomal dysfunction may constitute a common lipotoxic pathology across liver diseases. Ceramide synthases are sophisticatedly regulated, integrating hormonal, stress, and kinase signals. For instance, their transcription is controlled by factors like 17β-estradiol, GPER1, and FOXP3 (Qi et al., 2021; Wegner et al., 2014), and their activity is modulated post-translationally by phosphorylation via kinases such as CK2 and PKC (Fresques et al., 2015; Mullen et al., 2012). Our data position IL-6/JAK–STAT as a previously unrecognized upstream regulator of hepatic CERS, adding a critical inflammatory layer to this regulatory network. Notably, *CERS5* and *CERSC* emerge as actionable effectors of IL-6 associated hepatic dysfunction: higher *CERS5/C* expression correlates with reduced survival in hepatocellular carcinoma. A possible mechanistic basis is that these isoforms preferentially generate C16-ceramide, a species implicated in hepatic insulin resistance, steatosis, and inflammation (Chavez and Summers, 2012; Pewzner-Jung et al., 2010) and in the pathogenesis of obesity and type 2 diabetes (Raichur et al., 2019). Moreover, IL-6 has context-dependent roles in obesity and diabetes (Alexaki, 2024; Wueest and Konrad, 2020), suggesting that the IL-6/CERS axis may underlie a shared lipotoxic program when IL-6 drives liver dysregulation in these settings. Thus, whether hepatic autophagy is similarly impaired in these diseases warrants further investigation. Nevertheless, these observations nominate selective CERS5/6 inhibition as a rational strategy to mitigate lipotoxic stress and restore autophagy-lysosome competence in IL-6 related hepatic dysfunction.

In conclusion, we delineate a mechanistic link from chronic inflammation to failure of lipid homeostasis and lysosome-limited autophagy, identifying the IL-6/CERS axis as a key driver and therapeutic entry point of cancer-associated hepatic dysfunction.

## Acknowledgments

We thank Mujeeb Qadiri, Baolong Xia, Christians Villalta, Stephanie Mohr, Jonathan Zirin, Luping Liu, Litz Brown, and all members of the Perrimon Lab for their critical suggestions and help on this research. We thank Paula Montero Llopis and Microscopy Resources on the North Quad (MicRoN) core facility at Harvard Medical School for advice and help on confocal imaging. Young Yon Kwon, Yanshan Liang and Sheng Hui Lab for advice and help on lipidomics, and the *Drosophila* RNAi Screening Center (DRSC) and Bloomington Drosophila Stock Center (BDSC) for providing fly stocks used in this study. We thank Erika Bach (NYU School of Medicine) for the generous gift of Stat92E fly stocks. This work is funded in part by R01 DK136945, R01 AR057352, the Cancer Grand Challenges partnership funded by Cancer Research UK (CGCATF-2021/100022), and the National Cancer Institute (1 OT2 CA278685-01). Y.L. is supported by Charles A. King Trust. T. M. is supported by the Cystinosis Research Foundation and the Glenn Foundation for Medical Research Postdoctoral Fellowship in Aging Research. N.P. is an investigator of the Howard Hughes Medical Institute.

This article is subject to HHMI’s Immediate Access to Research policy, which requires that this article be made publicly available as initial and revised preprints deposited on a designated preprint server under a CC BY 4.0 license.

## Author Contributions

Y.L., T.M, and N.P. conceived the study and designed the experiments. Y.L. and T.M. conducted the majority of the experiments and the data analysis. Y.L. and A.W. carried out the mouse experiments. A.H.K performed AFM analysis. Y.L., T.M. and Y.H. were responsible for the bioinformatics analysis. J. M. A. helped for the lipidomics analysis. R.B generated the transgenic fly strains. Z.J.Z and X.M.S. helped for western blotting and plasmid construction. Y.L., T.M. and N.P. interpreted the results and wrote the manuscript with contributions and feedback from all authors. All authors reviewed and approved the final manuscript.

## Materials and Methods

### Drosophila strains

All flies were maintained on standard cornmeal fly food containing yeast and agar. Crosses were reared at 18°C to inhibit Gal4 and LexA. Adult offspring flies were collected 2 days after emerging, then kept at 18°C for an additional day and subsequently shifted to 29°C for indicated days to induce transgene and/or RNAi expression. Experimental fly food was changed every other day. Stocks used in this study include *esg-Le”A::GAD* (BDSC 66632), *tub-Gal80ts, Lpp-Gal4* (Song et al., 2017), *Le”Aop-Yki3SA-GFP* (Saavedra et al., 2021), *UAS-CG5966-RNAi* (RNAi #1: VDRC 13163, RNAi #2: VDRC 13164), *UAS-hop-RNAi* (BDSC 32966), *UAS-Stat92e-RNAi* (BDSC 33637), *UAS-schank-RNAi* (BDSC 29340), *UAS-Atg1-RNAi* (BDSC 26731), *UAS-HA-Stat92E* dominant-active form (Ekas et al., 2010), *UAS-Upd3* (Woodcock et al., 2015), and *UAS-Pvf1*(Xu et al., 2022). The *w1118* strain served as the control. Female flies are used in all experiments as they showed a more significant and consistent bloating phenotype.

### Plasmid and transgenic *Drosophila* strains

The full-length cDNA of *CG5966* (FlyBase ID: FBcl0163613) was obtained from DGRC (Stock 1514588). The insert was first cloned into pENTR-TOPO (Thermo Fisher) following the manufacturer’s protocol, then transferred into the *pWALIUM10-moe UAS* destination vector by Gateway LR recombination to generate *UAS-CG5966*. Positive clones were verified by colony PCR and complete Sanger sequencing. Endotoxin-reduced plasmid DNA was prepared by EndoFree midiprep for embryo injections. ΦC31 integrase–mediated transgenesis was used to integrate *UAS-CG5966* into standard attP docking sites: attP40 on chromosome 2L (25C7) or attP2 on chromosome 3L (68A4). Plasmid DNA was injected into pre-blastoderm embryos by in-house microinjection. Surviving G₀ adults were crossed individually to balancer stocks, and transformants were identified in the G₁ by *mini-white* marker selection. GAL4-dependent overexpression of *CG5966* was verified in the fat body by RT-qPCR.

### Lipidomics

Lipidomics and metabolomics of fly samples were performed using the classic Folch method (Folch et al., 1957). For each sample, 15 adult flies or 40 abdomens were used for lipidomics and metabolomics. Samples were homogenized in a 1 mL dounce homogenizer (TIGHT) on ice in 0.6 mL chloroform and 0.3 mL methanol (2:1 chloroform-methanol mixture). Homogenized samples were transferred to a 15 mL glass tube with a Teflon cap, with additional 0.6 mL chloroform and 0.3 mL methanol to make a final volume of 1.8 mL. Samples were vortexed briefly and incubated on a rotator for 30 minutes at room temperature. After incubation, 0.2 volumes (0.36 mL) of HPLC-grade deionized water (HPLC dH2O) were added. Samples were then vortexed three times for 5 seconds, followed by centrifugation 10 minutes at 1000 g at 4°C. Non-polar lipids solution (the lower phase) was transferred to a 1.85 mL glass tube using a glass pipette, then dried under nitrogen gas. Dried lipids pellets were stored at -80°C with nitrogen to prevent oxidation. Before injection, pellets were resuspended in 35 μL of HPLC-grade 50% methanol/50% isopropyl alcohol (IPA) and injected and analyzed using untargeted LC-MS/MS on a Thermo QExactive Plus Orbitrap mass spectrometer. Lipid molecules were identified using LipidSearch version 4.2 (Thermo Scientific). Ion intensity was quantified by measuring the area size of identified peaks.

### Protein lysate preparation and Western blot

30 abdomens with gonads and guts removed were dissected and lysed in RIPA lysis buffer (50 mM Tris, pH = 8.0, 150 mM NaCl, 0.1% SDS, 0.5% sodium deoxycholate, 1% Triton X-100, 1× complete ULTRA Mini EDTA-free protease inhibitor (Roche), and 1 mM PMSF). Dissected tissues in RIPA buffer were mixed with zirconium oxide beads (NextAdvance) and homogenized using a TissueLyser II (Qiagen). Homogenates were incubated on ice for 15 min and centrifuged for 15 min at 16,000 × g at 4 °C. Supernatants were transferred to new tubes, and protein concentrations were determined using the BCA Protein Assay Kit (Pierce) according to the manufacturer’s instructions. Samples were diluted to equal protein concentrations with RIPA buffer and stored at –80 °C. Equal amounts of total protein were mixed with 4× Laemmli sample buffer (Bio-Rad) containing 10% β-mercaptoethanol to a final 1× concentration, boiled at 95 °C for 5 min, and centrifuged at 16,000× g for 15 min at 4 °C. Supernatants were transferred to new tubes. Equal amounts of sample were loaded to a Mini-PROTEAN TGX Stain-Free Precast Gels (Bio-Rad) and ran using 1× Tris/Tricine/SDS Running Buffer (Bio-Rad) according to the manufacturer’s instructions. Spectra Multicolor Broad Range Protein Ladder (ThermoFish) was used as a molecular weight marker. The samples on the gel were transferred to a FL PVDF membrane using Tris/Glycine Buffer (Millipore) for 16 min, according to the manufacturer’s instructions. The membrane was then blocked with 1% Casein in 1× Tris Buffered Saline (TBS) (Bio-Rad) for 1 hour at room temperature. Next, the membrane was incubated in a primary antibody dilution in 1% Casein in 1× TBS overnight at 4 °C with gentle shaking. Primary antibodies were used at the following concentrations: 1:1000 mouse anti-Ctsl (RCD Systems, MAB22591), 1:2000 rabbit anti-Atg8 (Abcam, ab109364). Next, the membrane was washed three times quickly in TBST, followed by three 10-min washes in TBST with gentle shaking. The membrane was then incubated with fluorescently labeled secondary antibody in 1% Casein in 1× TBS for 1 h at room temperature with gentle shaking. The dilution of the fluorescently labeled secondary antibody was used as follow: 1:2500 hFAB™ Rhodamine; 1:2500 Goat anti-Rabbit IgG (H+L) Secondary Antibody, DyLight™ 800 4X PEG; 1:2500 IRDye 680RD Goat anti-Mouse IgG Secondary Antibody. After incubation, the membrane was washed three times quickly in TBST, followed by three 10-min washes in TBST with gentle shaking. Then, the membranes were imaged using a Biorad Chemidoc MP imager.

### Quantitative RT-PCR of fly samples

Nucleospin RNA kit (Macherey-Nagel) was used to extract RNA from fly samples according to the manufacturer’s protocol. cDNA was synthesized using iScript™ cDNA Synthesis Kit (Bio-Rad, 1708890) according to the manufacturer’s protocol. qPCR was performed with Thermal Cycler CFX 96 Real-Time System qPCR machine using iQ™ SYBR® Green Supermix (Bio-Rad). *RP49* and *CG13220* were used as housekeeping genes. qPCR primers used are:

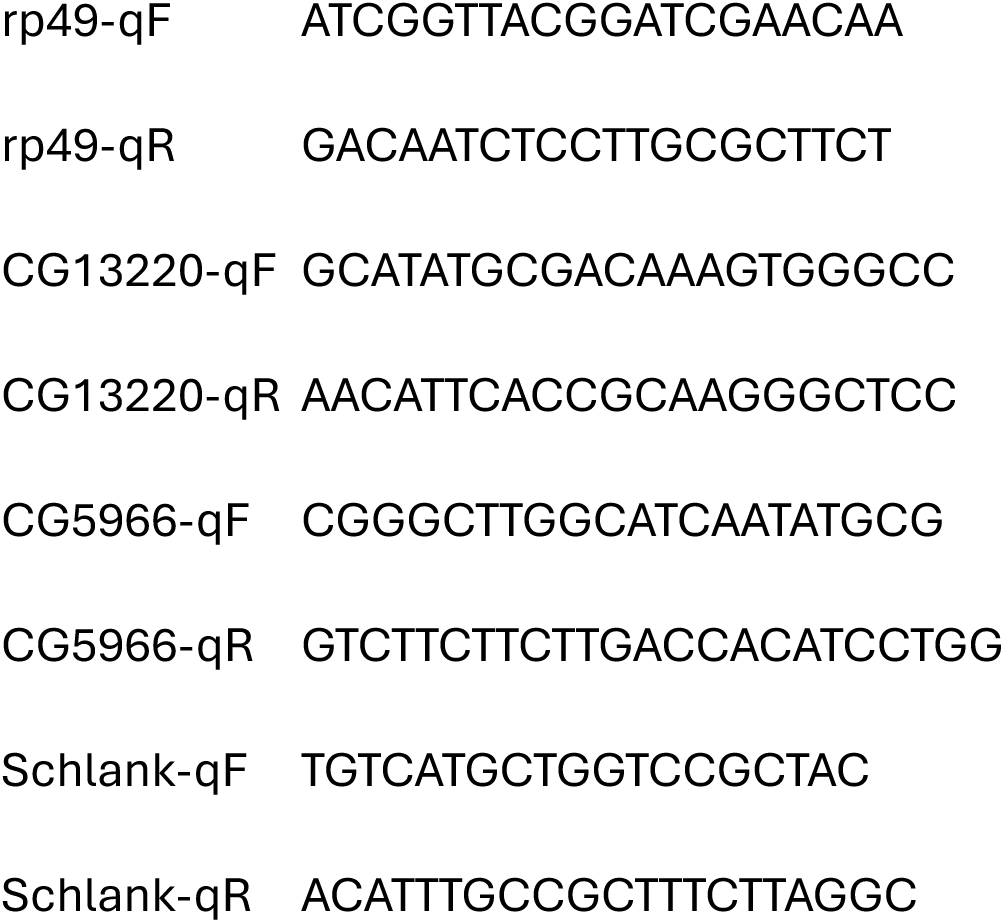

### Body fluid measurements

Body fluid measurement was followed a previous protocol (Liu et al., 2025). Briefly, flies were weighted before dried at 65 °C for 5 hours, and then weighted again. Body fluid is calculated by subtracting the second read from the first one.

### Climbing index and survival curve of flies

To assess the climbing ability, flies were transferred to a fresh vial and then tapped down to the bottom. Vials were imaged after 4 seconds. The percentages of flies in the upper 2/3 of the vial were recorded. The climbing index was generated from 3-4 independent vials for each genotype tested. Survival of flies was analyzed by calculating the percentage of flies alive in each vial, and 3 vials of 10-13 flies from each genotype were tested and flies were transferred to vials contain fresh food every other day or daily.

### Gut and fly imaging

Adult fly guts were dissected in cold PBS and immediately fixed for 30 minutes in 1X PBS contain 4% formaldehyde. Guts were washed three times with 1X PBS contain 0.3% Triton X-100 (PBT) and then mounted in Vectashield with DAPI (Vector Laboratories, H-1200). For pH3 staining, washed guts were incubated with blocking buffer (5% NDS in 1X PBS contain 0.3% Triton X-100) for 2 hours at room temperature, incubated with rabbit anti-pH3 (1:500, CST 9701S) antibody overnight at 4°C. Guts were washed three times in PBT next day before incubating with secondary antibody, donkey anti-rabbit 565 (1:2000, Molecular Probes A31572), for 1 hour. After three final washes with PBT, guts were mounted in Vectashield with DAPI (Vector Laboratories, H-1200). Confocal images were taken with Nikon Ti2 Spinning Disk with Nikon Elements Acquisition Software AR (v5.02). Tumors were quantified by measuring the average GFP signaling strength of gut tumors from individual flies using Fiji-imageJ. Adult fly phenotypes were imaged using a ZEISS Axiozoom V16 fluorescence microscope. Bloating was quantified by the ratio between abdomen and head compartment size using Fiji-imageJ.

### Live imaging of autophagic vesicle and lysosome

For live imaging of autophagy and lysosomes, abdominal cuticles were dissected in cold Schneider’s insect medium. Samples were then incubated for 10 minutes at room temperature in a staining buffer composed of Schneider’s medium, CYTO-ID® Autophagy Detection Reagent (1:1000, Enzo Life Sciences ENZ-KIT175), Hoechst nuclear stain (1:1000), and LysoTracker™ Deep Red (500 nM, Thermo Fisher Scientific L12492). Following incubation, the samples were rinsed once with fresh Schneider’s medium and then twice with phosphate-buffered saline (PBS). Finally, the stained cuticles were transferred to a glass slide with mounting medium, coverslipped, and immediately visualized using a Nikon Ti2 Spinning Disk confocal microscope.

### Chromatin Immunoprecipitation

ChIP assay was performed with the SimpleChIP® Plus Enzymatic Chromatin IP Kit (Cell Signaling, 9005). Fat bodies from 50 adult flies were collected for each sample. Fat body expressing HA-tagged dominant-active Stat92e were dissected and flash-frozen in liquid nitrogen. Cross-linked was done with 1.5% formaldehyde for 20 minutes at room temperature. Cross-linking was stopped by adding glycine solution for 5 minutes at RT, samples were washed twice with 1 ml 1X PBS containing 1X Protease Inhibitor Cocktail and homogenized using 1 ml dounce homogenizer to release nuclei. Samples were lysed with Diagenode Bioruptor sonicator to retrieve cross-linked chromatin. Chromatins were diluted in 1X ChIP buffer before incubating with 10 ul HA-Tag (C29F4) Rabbit mAb (Cell Signaling, 3724) or Normal Rabbit IgG (Cell Signaling, 2729) overnight with rotation at 4°C. Next day, each immunoprecipitation was incubated for 2 hours with 30 ul ChIP-Grade Protein G Magnetic Beads (Cell Signaling, 9006) at 4 °C with rotation. Beads were then washed and incubated in 150 µl 1X ChIP Elution Buffer at 65 °C for 30 minutes with vortexing (1200 rpm) to elute the chromatin. 6 µl 5M NaCl and 2 µl Proteinase K was added to the eluted chromatin supernatant to reverse cross-links and incubated 2 hours at 65°C. DNA was then purified from each sample using Spin Columns provided by the kit. 1 ul DNA sample was used as template for qPCR to detect enrichments of certain DNA regions. qPCR of a fragment in the Sam-S gene region was used as the negative control. Primers used are:

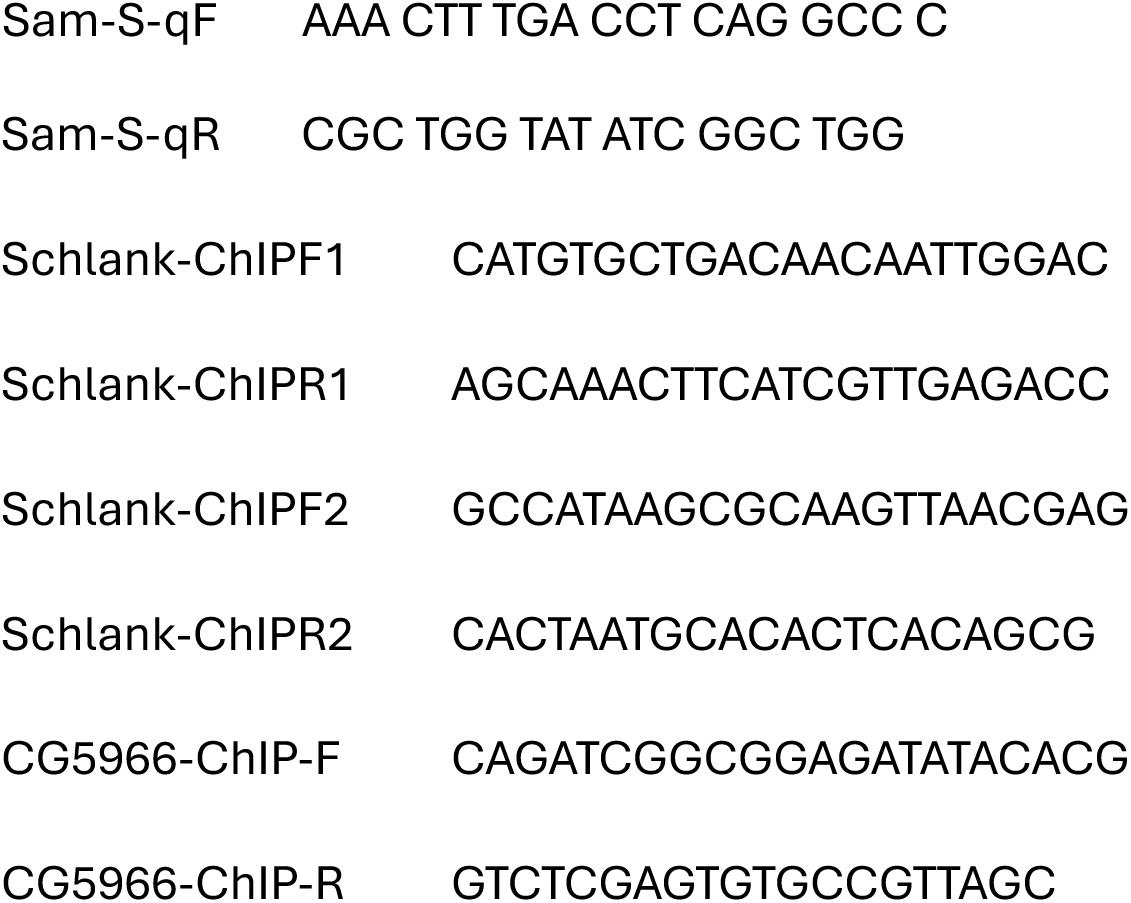

### Mouse models

The *Kras^G12D/+^*; *Lkb1^f/f^* mice described before (Ji et al., 2007) were further backcrossed to FVB mice. Tumor induction in adult 12- to 20-week-old FVB mice was achieved by intranasal administration of 75 μL of PBS containing 2.5 × 10^7^ pfu of Adenovirus CMV-Cre (Ad5CMV-Cre) obtained from the University of Iowa Gene Transfer Vector Core (Iowa City, IA) and 1 mM CaCl_2_. Male mice were used for liver RNAseq. The *Kras^G12D/+^*; *Lkb1^f/f^* mice experiments were approved by the Institutional Animal Care and Use Committee (IACUC) of Weill Cornell Medical College and maintained as approved by the Institutional Animal Care and Use Committee (IACUC) at Weill Cornell Medicine (NYC).

C57BL/6J mice were obtained from the Jackson Laboratory (Strain #000664). 2 x 10^6^ Lewis Lung Carcinoma (LLC) cells or LLC cells edited to produce IL-6 (LLC-IL6) were subcutaneously inoculated into their right flank after a week of acclimation. Mice were kept in pathogen-free conditions on a 24 hour 12:12 light-dark cycle. 9 weeks old male C57BL/6J mice were used in the experiments. C57BL/6J mice experiments were approved by the Institutional Animal Care and Use Committee (IACUC) at Cold Spring Harbor Laboratory (CSHL) and were conducted in accordance with the National Institutes of Health Guide for the Care and Use of Laboratory Animals. Body weights and clinical signs of cachexia were monitored daily. Handling was kept to a minimum. Mice were sacrificed when tumor size exceeded 2 cm length, when weight loss exceeded 15% from peak weight, or when showing clinical signs of discomfort indicative of cachectic endpoint as stated by the Animal Cachexia Score (ACASCO): piloerection, diarrhea or constipation, hunched posture, tremors, and closed eyes. Death was confirmed by cervical dislocation.

### Lewis Lung Carcinoma (LLC) cell line

LLC cells were purchased from American Type Culture Collection (ATCC), cultured in complete growth medium consisting of Dulbecco’s Modified Eagle Medium (DMEM) (#10027CV; Corning) containing 1x Penicillin-Streptomycin solution (#15-140-122; Thermo Fisher) and 10% of Heat-Inactivated Fetal Bovine Serum (FBS) (#10-438-026; Thermo Fisher). LLC cells were transfected with 500ng of plasmid (comprising 2.5:1 PB-IL6 plasmid and PBase plasmids) using Lipofectamine 3000 (Thermo Fisher) according to the manufacturer’s protocol. PBase plasmid was obtained from System Biosciences (#PB210PA-1) and PB-IL6 was obtained from VectorBuilding comprising mouse Il6 cDNA driven by EF1a promoter and flanked by piggyBac elements. 48 hours after transfection, the media was changed and replaced with DMEM media supplemented with 3 μg/ml puromycin. After 14 days of antibiotic selection, monoclonal populations were isolated by serial dilutions in a 96-well plate. To identify clones with constitutive IL-6 expression, we measured IL-6 in the cell supernatant for each clone using the Mouse IL-6 ELISA Kit (#ab222503; Abcam).1x Trypsin-EDTA (#15400054; Thermo Fisher) was used for cell dissociation. Cells were resuspended in FBS-free DMEM and counted using a Vi-Cell counter prior to subcutaneous injection. For each C57BL/6J mouse, 2×10^6^ viable cells diluted in 100μL DMEM into the right flank.

### Lipid extraction and profiling in mice

Lipid extracts were analyzed using a Dionex Ultimate 3000 RSLC system (Thermo Scientific) coupled with a QExactive mass spectrometer (Thermo Scientific, Waltham, MA, USA) mass spectrometer. Chromatographic separation was achieved on an ACQUITY UPLC CSH C18 column (130Å, 1.7 µm, 2.1 mm × 100 mm) with an ACQUITY UPLC CSH C18 VanGuard pre-column (130Å, 1.7 µm, 2.1 mm × 5 mm) (Waters, Milford, MA) with column temperature at 50 °C. For the gradient, mobile phase A consisted of an acetonitrile-water mixture (6:4), and mobile phase B was a 2-propanol-acetonitrile mixture (9:1), both phases containing 10 mM ammonium formate and 0.1% formic acid. The linear elution gradient was: 0-3 min, 20% B; 3-7 min, 20-55% B; 7-15 min, 55-65% B; 15-21 min, 65-70% B; 21-24 min, 70-100% B; and 24-26 min, 100% B, 26-28 min, 100-20% B, 28-30 min, 20% B, with a flow rate of 0.35 mL/ min. The autosampler was at 4°C. The injection volume was 5 µL. Needle wash was applied between samples using a mixture of dichloromethane-isopropanol-acetonitrile (1:1:1).

ESI-MS analysis was performed in positive and negative ionization polarities using a combined full mass scan and data-dependent MS/MS (Top 10) (Full MS/dd-MS^2^) approach. The experimental conditions for full scanning were as follows: resolving power, 70,000; automatic gain control (AGC) target, 1 × 10^6^; and maximum injection time (IT), 100 ms. The scan range of the instrument was set to m/z 100-1200 in both positive and negative ion modes. The experimental conditions for the data-dependent product ion scanning were as follows: resolving power, 17,500; AGC target, 5 × 10^4^; and maximum IT, 50 ms. The isolation width and stepped normalized collision energy (NCE) were set to 1.0 m/z, and 10, 20, and 40 eV. The intensity threshold of precursor ions for dd-MS2 analysis and the dynamic exclusion were set to 1.6 × 10^5^ and 10 s. The ionization conditions in the positive mode were as follows: sheath gas flow rate, 50 arb; auxiliary (AUX) gas flow rate, 15 arb; sweep gas flow rate, 1 arb; ion spray voltage, 3.5 kV; AUX gas heater temperature, 325 ^◦^C; capillary temperature, 350 ^◦^C; and S-lens RF level, 55. The ionization conditions in the negative mode were as follows: sheath gas flow rate, 45 arb; auxiliary (AUX) gas flow rate, 10 arb; sweep gas flow rate, 1 arb; ion spray voltage, 2.5 kV; AUX gas heater temperature, 320 ^◦^C; capillary temperature, 320 ^◦^C; and S-lens RF level, 55.

Thermo Scientific™ LipidSearch™ software version 5.0 was used for lipid identification and quantitation. First, the product search mode was used during which lipids were identified based on the exact mass of the precursor ions and the mass spectra resulting from product ion scanning. The precursor and product tolerances were set to 10 and 10 ppm mass windows. The absolute intensity threshold of precursor ions and the relative intensity threshold of product ions were set to 30000 and 1%. Next, the search results from the individual positive or negative ion files from each sample were aligned within a retention time window (±0.25 min) and then all the data were merged for each annotated lipid with a retention time correction tolerance of 0.5 min. The annotated lipids were then filtered to reduce false positives by only including a total grade of A or B or C for PC and SM; otherwise only including grade A or B.

### Bulk RNA-Sequencing from KL livers

Total RNA was extracted from the liver using TRIzol (Thermo Fisher), followed by a clean-up using RNeasy kit (Qiagen). 1mg RNA from each sample was submitted to the WCM Genomics Resources Core Facility. Raw sequenced reads were aligned to the mouse reference GRCm38 using STAR (v2.4.1d, 2-pass mode) aligner, and raw counts were obtained using HTSeq (v0.6.1). Differential expression analysis, batch correction and principal component analysis (PCA) were performed using R Studio Version 4.2.2 and DESeq2 (v.1.38.3). Gene set enrichment analysis (GSEA) analysis was performed with the R package fGSEA (<u>10.18129/B9.bioc.fgsea</u>), using the Reactome pathway database contained in the 2022 release of Mouse Molecular Signatures Database from the Broad Institute (https://www.gsea-msigdb.org/gsea/msigdb/mouse/collections.jsp).

### Quantitative RT-PCR of mouse samples

For LLC and IL6-secreting LLC mouse liver samples, 100 mg tissue was lysed in 1 mL of Qia-zol with Qiagen TissueLyser II at 20 Hz for 4 minutes. Samples were centrifuged at 12,000 g for 12 minutes at 4 C to separate aqueous layer from others. Approximately 600 ul of RNA containing aqueous layer was collected and processed using Qiagen RNeasy lipid RNA extraction kit and QiaCube machine. RNA were diluted in ddH20 to 100 ng/uL. cDNA was synthesized using iScript™ cDNA Synthesis Kit (Bio-Rad, 1708890) according to the manufacturer’s protocol. qPCR was performed with Thermal Cycler CFX 96 Real-Time System qPCR machine using iQ™ SYBR® Green Supermix (Bio-Rad). RN18s was used as housekeeping gene. Primers used are:

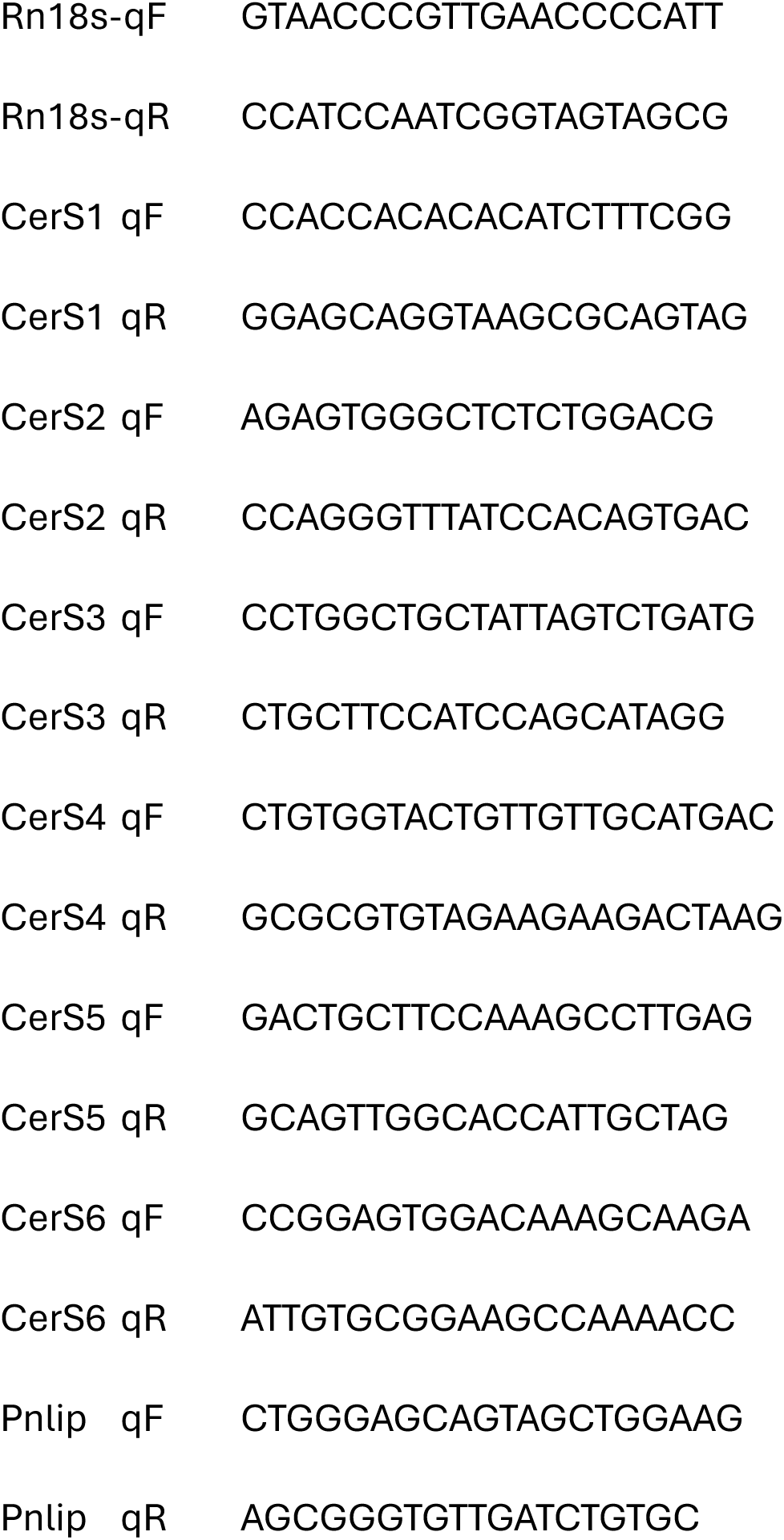
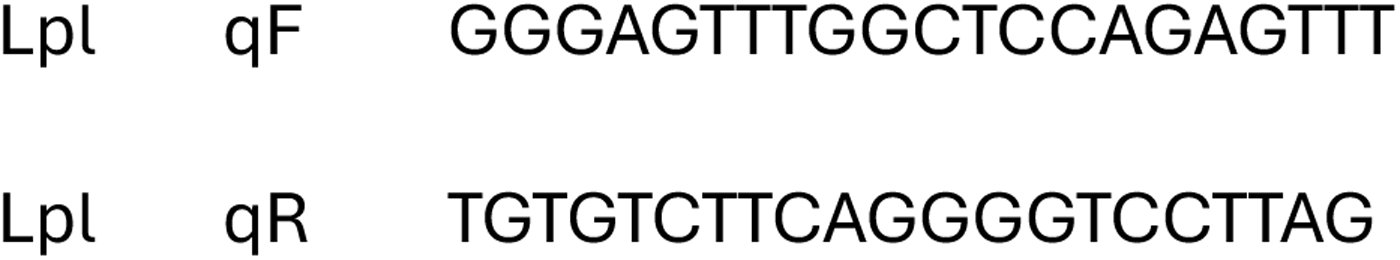

### Public datasets, Software, Quantification and Statistical Analyses

Transcriptomic data from non-diseased liver samples of 226 individuals were obtained from the Genotype-Tissue Expression (GTEx) Project. Transcriptomic data and corresponding survival records were obtained from the Hepatocellular Carcinoma (LIHC) cohort of 372 patients in the PanCancer Atlas of The Cancer Genome Atlas (TCGA) Program (Hoadley et al., 2018; Liu et al., 2018). Sex and gender were not considered in the design of present research, and related information was not collected for the analysis. Kaplan-Meier survival curves were generated using the R packages survival and survminer. We used Biorender and PyMOL for a subset of figures, and ChatGPT and Gemini were used to proofread the text. The authors reviewed and edited the content as needed and take full responsibility for the content of the publication. GraphPad Prism was used for generation of figures, statistical analysis, and gene expression correlation analysis. Data distribution was assumed to be normal, but this was not formally tested. Animals and samples were randomly assigned to experimental groups, and the data collection was randomized. Data collection and analysis were not performed blind to the conditions of the experiments. Statistical analysis was done with student’s t test (two experimental groups) or ordinary one-way ANOVA (three or more experimental groups) by the default settings of GraphPad Prism (ns indicates not significant, ∗ indicates p<0.05, ∗∗ indicates p<0.01, ∗∗∗ indicates p<0.001, ∗∗∗∗ indicates p<0.0001). Gene expression levels (qPCR) were normalized to the mean of control samples. Error bars indicate the standard deviations with the mean as the center. n indicates the number of biological replicates in each experiment, for fly experiments, each biological replicates contains 10-16 flies otherwise noted; for mouse experiment, each biological replicates is one mouse.

---

**Figure S1, related to Figure 1.**
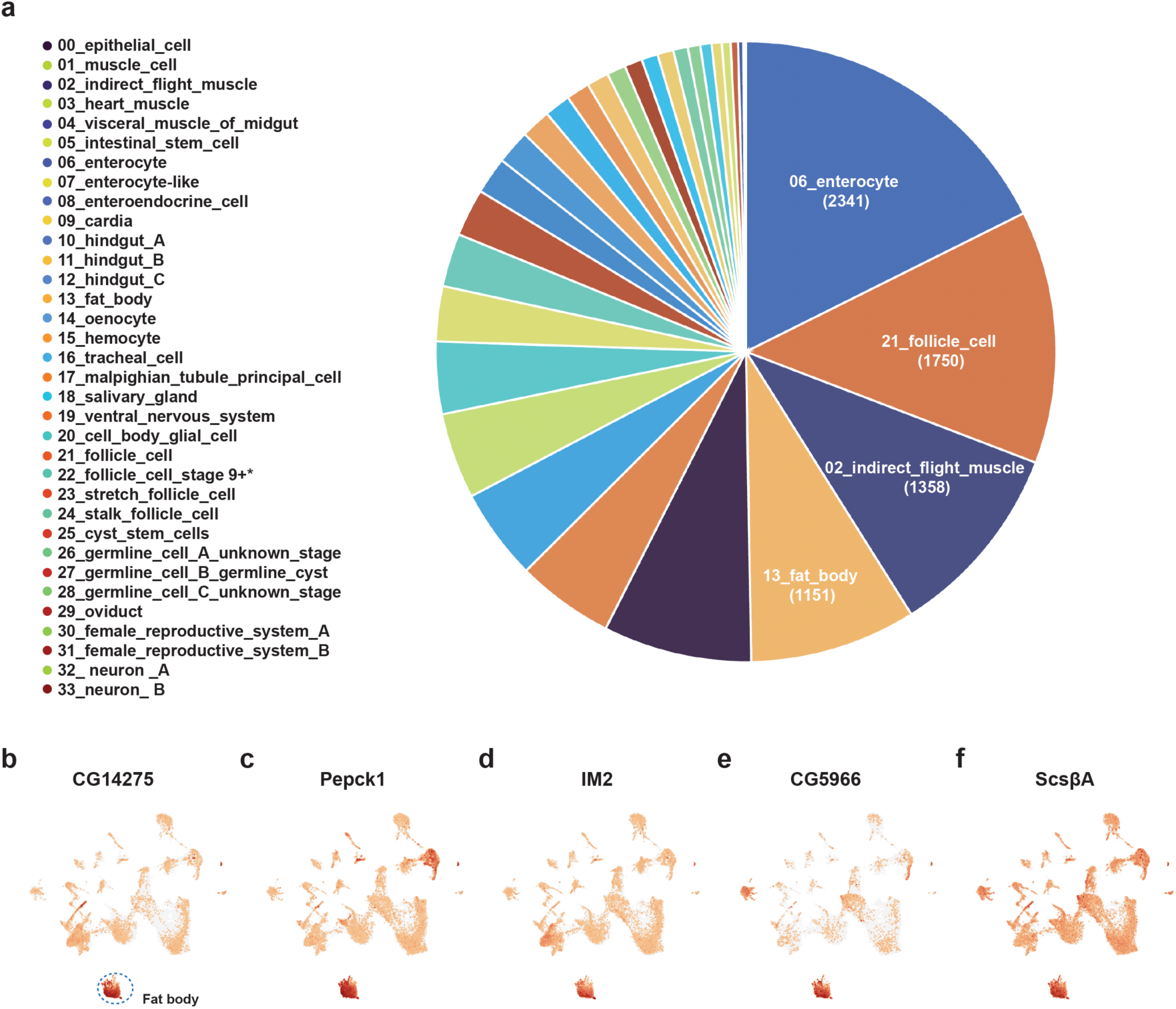
**a**, Distribution of progression associated genes across cell clusters. **b**-**f**, Expression of CG14275 (**b**), Pepck1 (**c**), IM2 (**d**), CG5966 (**e**), ScsbetaA (**f**) across cell clusters, data retried from snRNA-seq.

**Figure S2, related to Figure 2.**
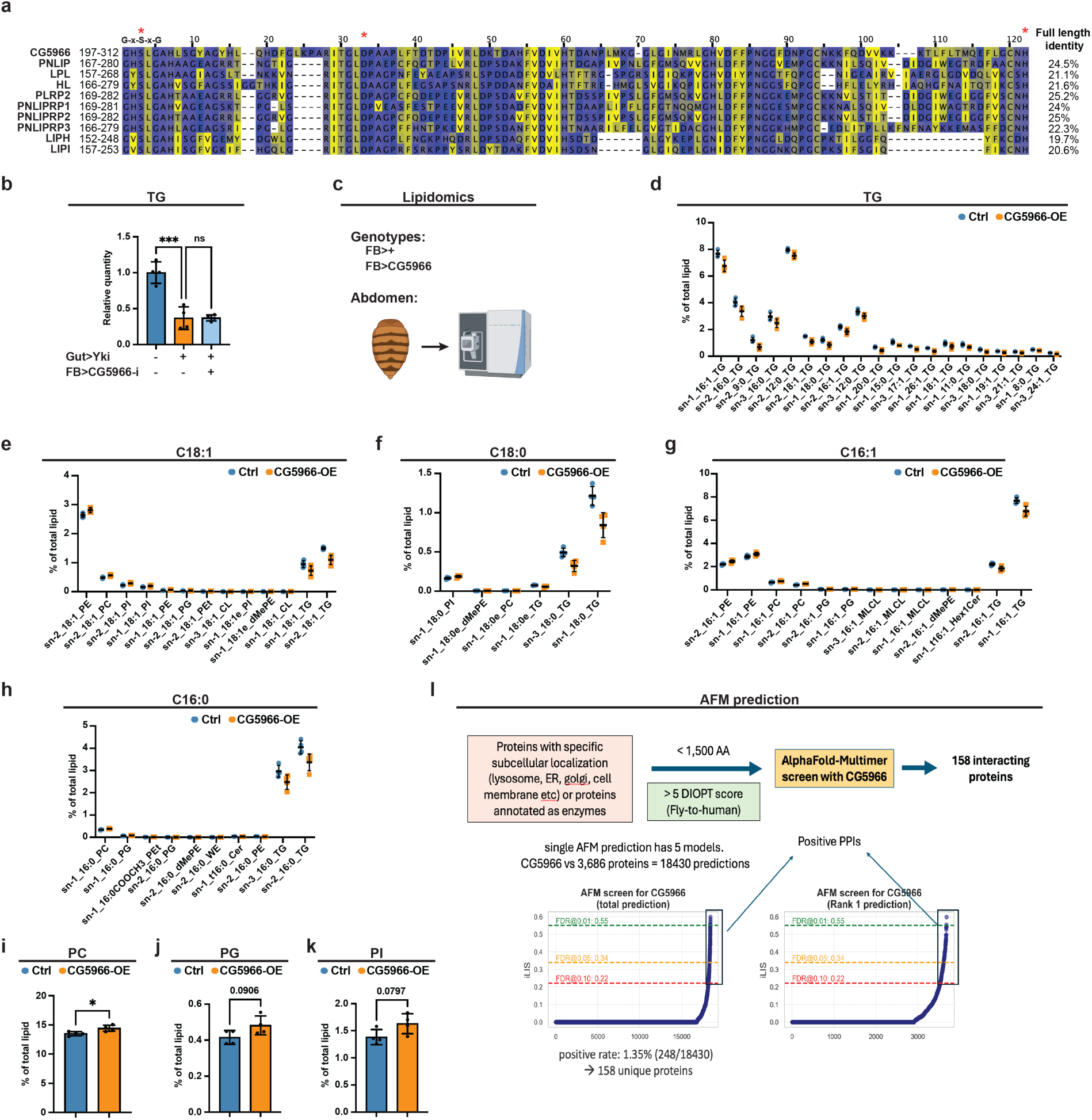
**a**, Amino acid sequence alignments showing similarity of human lipases to *CG5966*. **b**, Whole-body TG levels of Yki flies with or without fat-body *CG5966* depletion at day 6 (n = 4). **c**, Lipidomics analysis setting. **d**-**h**, Proportions of significantly changed abdomen TG species (**d**), and lipid species containing C18:1 (**e**), C18:0 (**f**), C16:1 (**g**), or C16:0 (**h**) acyl chains in flies with or without fat-body *CG5966* overexpression (n=4), data were retrieved from lipidomics analysis. **i**-**k**, Proportions of abdomen PC (**i**), PG (**j**), PI (**k**) of flies with or without fat-body CG5966 overexpression (n=4), data were retrieved from lipidomics analysis. **l**, AFM analysis setting. *p < 0.05, **p < 0.01, ***p < 0.001, ****p < 0.0001, ns indicates not significant. Error bars indicate SDs. n indicates the number of biological replicates in each experiment.

**Figure S3, related to Figure 3.**
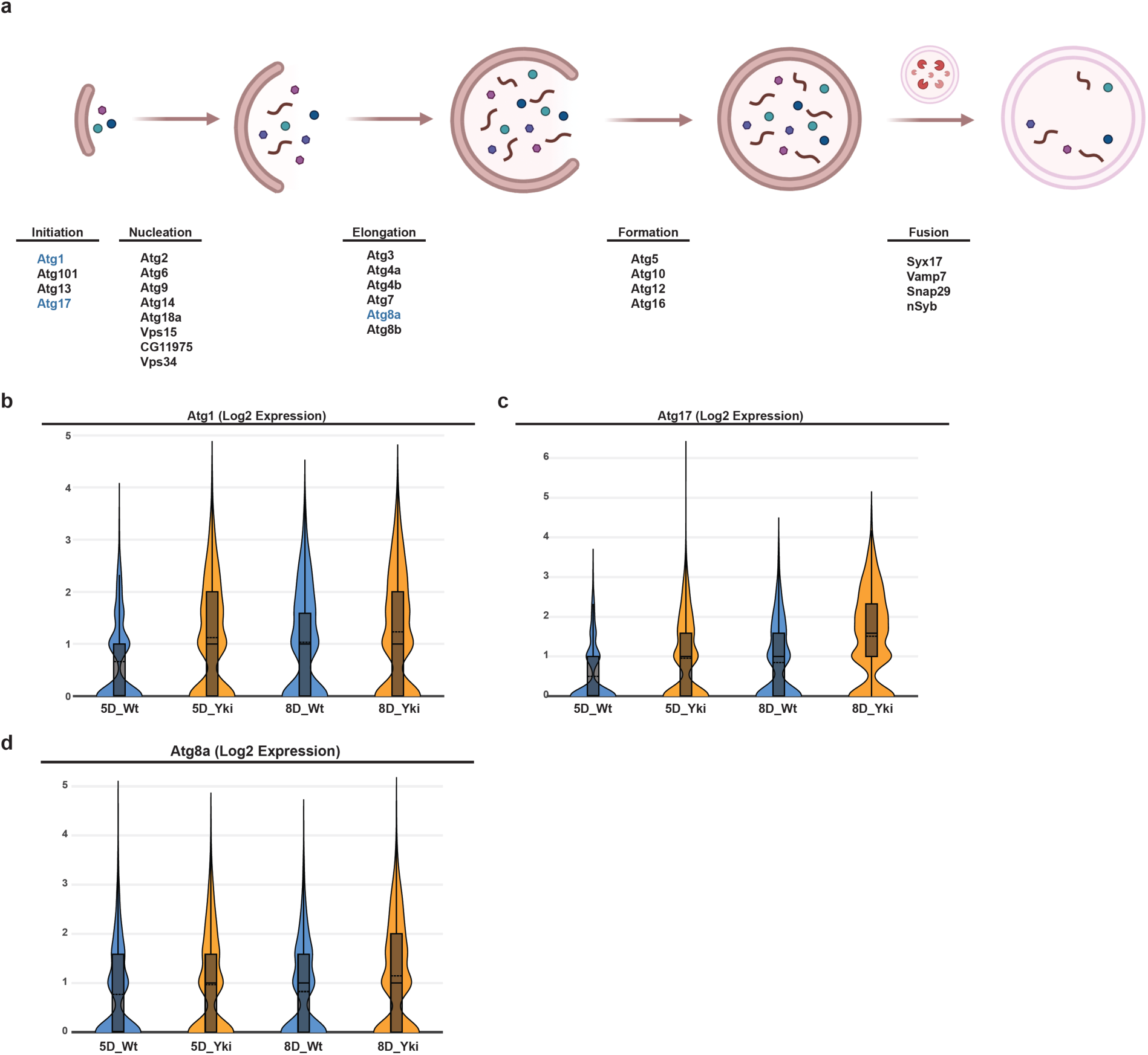
**a**, A schematic of autophagy in *Drosophila*, genes upregulated in Yki are labeled in blue, other genes are not significantly changed. **b-d**, Log2 expression of *Atg1* (**b**), *Atg17* (**c**), and *Atg8a* (**d**) in day 5 and day 8 control and Yki flies, data were retrieved from snRNAseq analysis.

**Figure S4, related to Figure 4.**
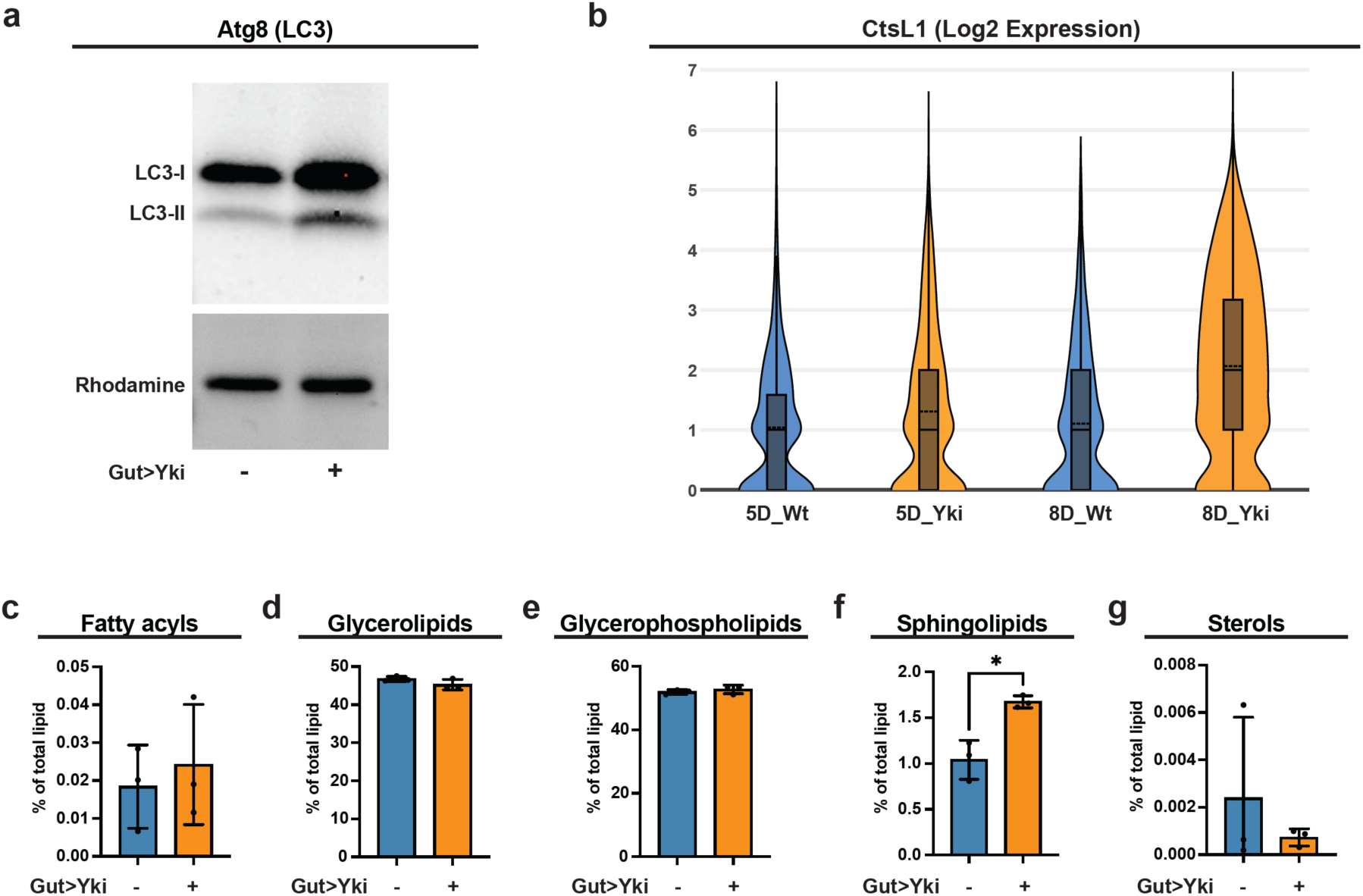
**a**, Western blot of Atg8 (LC3) of abdomen samples from control and Yki flies. Rhodamine was used as loading control. **b**, Log2 expression of *CtsL1* in day 5 and day 8 control and Yki flies, data were retrieved from snRNAseq analysis. **c**-**g**, Whole-body proportions of lipid subgroups fatty acyls (**c**), glycerolipids (**d**), glycerophospholipids (**e**), sphingolipids (**f**), and sterols (**g**) in control and Yki flies, data were retrieved from lipidomics analysis.

**Figure S5, related to Figure 5.**
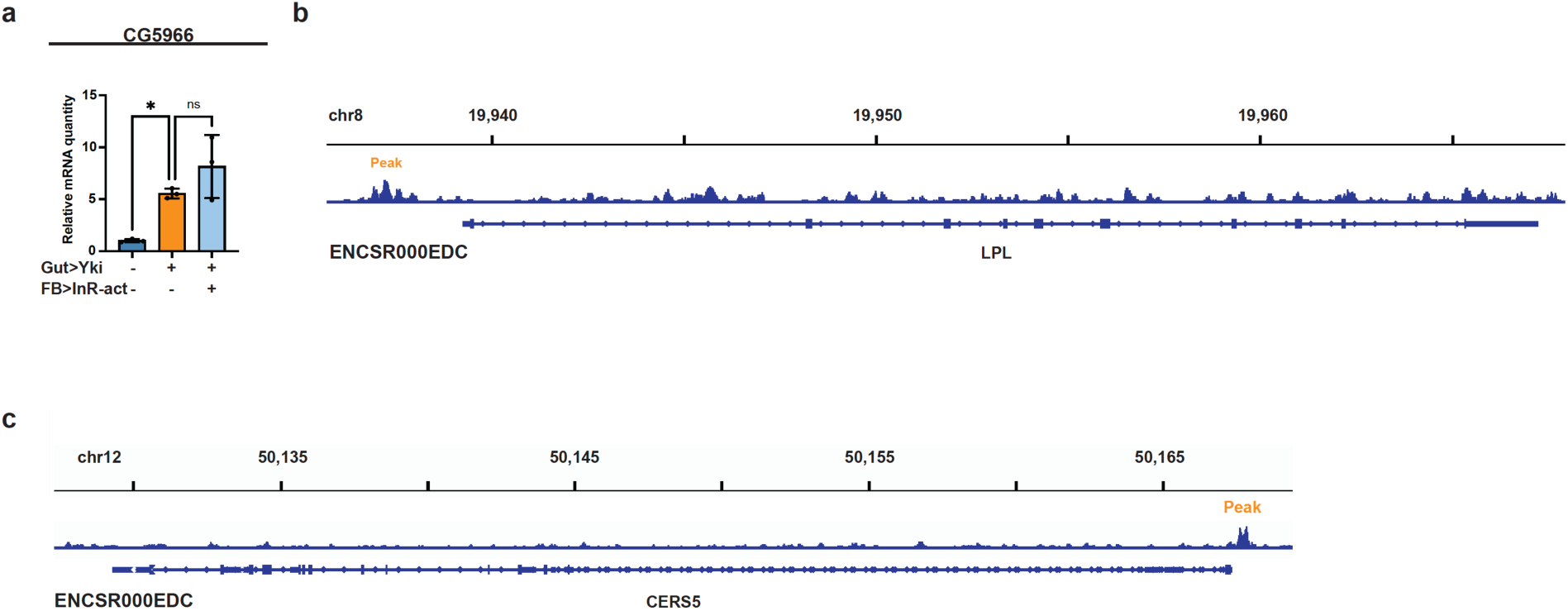
**a**, *CG5SCC* mRNA expression in the fat body of control, Yki flies, and Yki flies with fat-body *InR-act* (active form of InR) expression at day 6 (n = 3). *p < 0.05, **p < 0.01, ***p < 0.001, ****p < 0.0001, ns indicates not significant. Error bars indicate SDs. n indicates biological replicates. **b**,**c**, Data from the ChIP-seq data (ENCSR000EDC) indicating enrichment of STAT3 binding at the *LPL* (**b**) and *CERS5* (**c**) promoter and gene regions.

**Figure S6. related to Figure 6.**
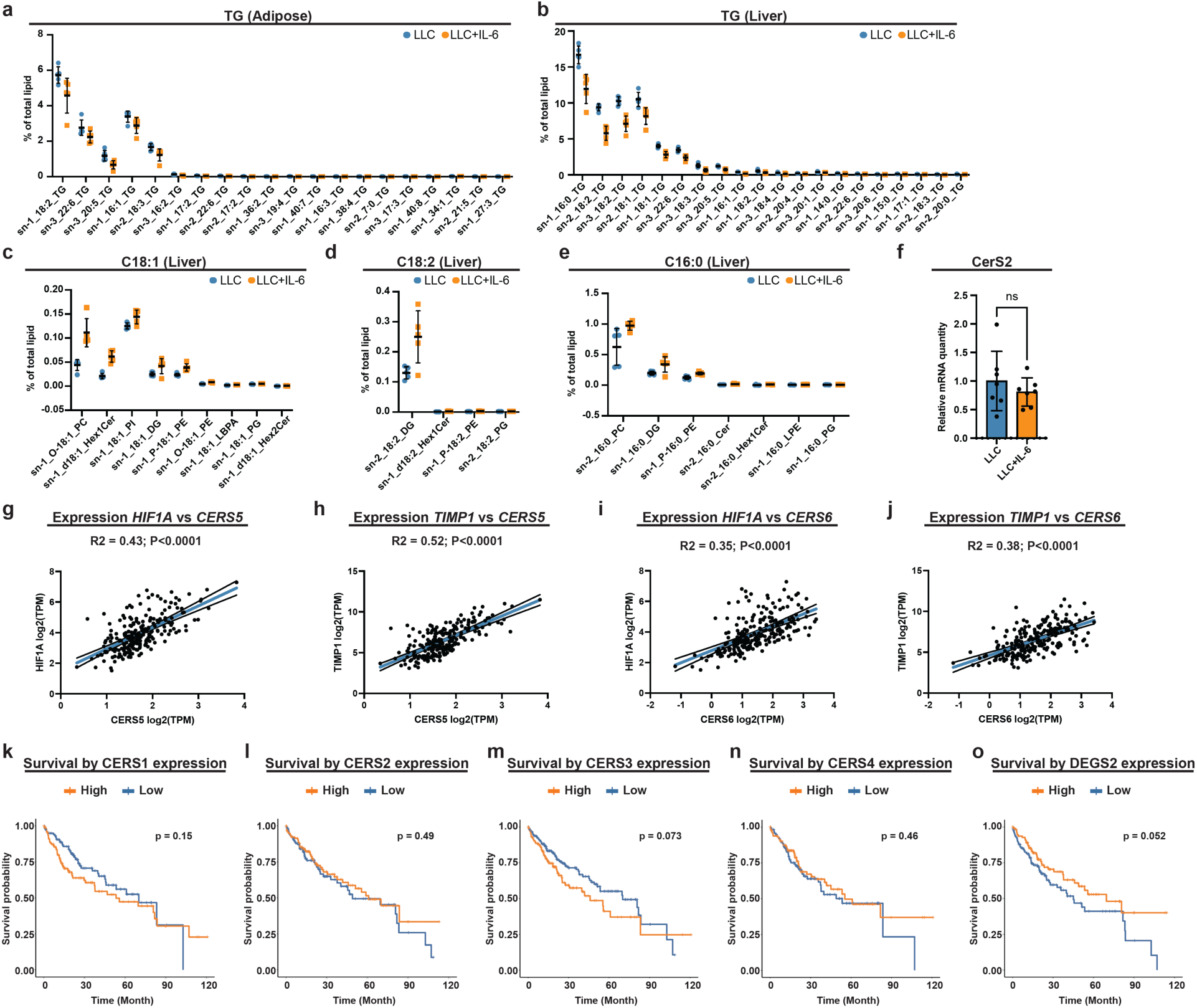
**a**,**b**, Proportions of significantly changed adipose (**a**) and liver (**b**) TG levels of B6 mice injected with LLC cells with or without IL-6 expression (n=5). **c**-**e**, Proportions of significantly changed lipid species containing C18:1 (**c**), C18:2 (**d**), or C16:0 (**e**) acyl chains of B6 mice injected with LLC cells with or without IL-6 expression in liver (n=5). **f**, qRT–PCR of *CerS2* mRNA in liver of B6 mice injected with LLC cells with or without IL-6 expression (n = 7). **g**-**j**, Correlation plots showing the positive relationship between expression of *HIF1A* and *CerS5* (**g**), *TIMP1* and *CerS5* (**h**), *HIF1A* and *CerS6* (**i**), *TIMP1* and *CerS6* (**j**) in non-diseased liver samples of 226 participants in the GTEx Project. TPM, transcript per million. **k**-**o**, Kaplan–Meier survival curves displaying the estimated survival probabilities of people with hepatocellular carcinoma with low (bottom third) and high (top third) hepatic expression of *CERS1* (**k**), *CERS2* (**l**), *CERS3* (**m**), *CERS4* (**n**), and *DEGS2* (**o**). *p < 0.05, **p < 0.01, ***p < 0.001, ****p < 0.0001, ns indicates not significant. Error bars indicate SDs. n indicates the number of biological replicates in each experiment.

## Notes

### Competing Interest Statement

The authors have declared no competing interest.

### Summary of Updates

There has been substantial revision to the manuscript

